# The chromatin factor ROW cooperates with BEAF-32 in regulating long-range inducible genes

**DOI:** 10.1101/2021.03.08.434270

**Authors:** Neta Herman, Sebastian Kadener, Sagiv Shifman

## Abstract

Insulator proteins located at the boundaries of topological associated domains (TAD) are involved in regulating chromatin loops. Yet, how chromatin loops contribute to transcription regulation is still not clear. Here we show that Relative-of-WOC (ROW) is essential for the long-range transcription regulation mediated by the Boundary Element-Associated Factor of 32kD (BEAF-32). We found that ROW physically interacts with heterochromatin proteins (HP1b and HP1c) and the insulator protein BEAF-32. The co-localization happens at TAD boundaries where ROW, through its AT-hooks motifs, binds AT-rich sequences flanked by BEAF-32 binding sites and motifs. Knockdown of *row* resulted in downregulation of genes that are long-range targets of BEAF-32 and bound indirectly by ROW (without binding motif). Analysis of high-throughput chromosome conformation capture (Hi-C) data revealed long-range interactions between promoters of housekeeping genes bound directly by ROW and promoters of developmental genes bound indirectly by ROW. Thus, our results show cooperation between BEAF-32 and the ROW complex, which includes HP1 proteins, to regulate the transcription of developmental and inducible genes by chromatin loops.

## Introduction

Chromosomes are organized at the sub-megabase scale into domains with a high level of interactions within them, known as topologically associating domains (TADs), separated by sharp boundaries with a lower level of interactions between domains (1, 2). TADs are frequently linked with a specific chromatin state and can be divided into active, PcG, HP1, and inactive TADs (2). Most TAD boundaries in *Drosophila* are located at active promoters; however, it is not clear whether this genomic distribution regulates transcription or is a result of transcriptional activity (3).

The *Drosophila* insulator protein BEAF-32 is bound to TAD boundaries located at promoters of housekeeping genes (4). Those promoters are characterized by multiple BEAF-32 binding motifs surrounded by AT-rich spacers (5). BEAF-32 has been implicated in both transcription and genome organization. For example, knockout of BEAF-32 causes defects in the morphology of the male X polytene chromosome, suggesting that it is involved in chromatin structure and or dynamics (6). It is still not clear what is the role of BEAF-32 in TAD boundaries formation as one study showed that depletion of *BEAF-32* does not affect TAD boundaries (7), while a more recent study showed that it does change both TAD boundaries and loops (8). BEAF-32 is associated with transcription regulation through multiple mechanisms, including regulation of enhancer-promoter interactions (9), through long-range contacts between promoters bound directly and indirectly by BEAF-32 (10, 11), and by restricting the deposition of H3K9me3, H3K27me3, and the spread of heterochromatin to actively transcribed promoters (5, 11).

A key component in the formation and maintenance of heterochromatin is the protein HP1 – a highly conserved protein first identified in *Drosophila* (now called HP1a) (12, 13). *Drosophila* has five HP1 paralogs, three that are ubiquitously expressed (HP1a, HP1b, and HP1c) and two which are specific for the germline (HP1d and HP1e) (14). Most of what we know about the HP1 family is from studies focused on *Drosophila* HP1a. HP1a is involved mostly in heterochromatin formation and transcription silencing and bind to H3K9me2/3 (15, 16). The potential functions of HP1b and HP1c are less explored. HP1b is distributed at both euchromatin and heterochromatin, whereas HP1c was found mainly at euchromatin (17–21). Although HP1c can bind H3K9me2/3 *in vitro*, it was shown to be localized at active promoters with poised RNA polymerase II (RNA pol II) (20). The molecular mechanisms that determine the different genomic distribution of the HP1 paralogs remain mostly unknown.

The *Drosophila* HP1c interacts with two zinc finger proteins, Without Children (WOC) and Relative Of WOC (ROW), as well as with the ubiquitin receptor protein Dsk2 (20, 22, 23). Recently it was suggested that the localization of the HP1c complex at euchromatin is dependent on ROW (24). ROW contains protein domains that implicate it in transcription regulation, including multiple zinc-finger (ZNF) motifs, AT-hooks, and a glutamine-rich domain in the C-terminal, that resemble activation domains found in transcription factors (20). Strikingly, the knockdown of *row*, *woc,* and *HP1c* leads to expression changes of a common set of genes (20). Moreover, several lines of evidence suggest that the complex containing ROW, WOC, and HP1c is involved in transcription activation. First, HP1c interacts with the Facilitates Chromatin Transcription Complex (FACT) to recruit FACT to active genes and the active form of RNA pol II (25). Second, HP1c interacts with the ubiquitin receptor protein Dsk2, which is involved in the positive regulation of transcription (22). Third, results of chromatin immunoprecipitation in *Drosophila* S2 cells followed by high-throughput sequencing (ChIP-seq) of ROW, WOC, and HP1c revealed localization of the complex around the transcription start sites (TSSs) of actively transcribed genes (22). Finally, depletion of *row* in S2 cells leads to the downregulation of approximately 80% of the genes that are both differentially expressed and are targets of the complex (22). However, *in vivo* expression analysis with RNAi lines of *row*, *woc,* and *HP1c* (RNA from whole larvae) showed similar numbers of upregulated and downregulated genes (20).

ROW is also an ortholog of *POGZ* – a human risk gene for neurodevelopmental disorders (26) that interacts with heterochromatin proteins (27). Although *row* is expressed throughout most *Drosophila* tissues and developmental stages, it displays the highest expression level in the larval central nervous system (26). Moreover, neuron-specific knockdown of *row* in adult flies affects non-associative learning (26). Thus, *row* is similar to *POGZ* as both interact with heterochromatin proteins and are involved in neurodevelopment and learning (28).

Here, we comprehensively characterized *row* and its binding partners for the first time *in-vivo* in adult *Drosophila*. We found that knockdown of *row* using constitutive promoter results in reduced viability, fertility, and changes in the expression of metabolic-related genes. Interestingly, we found that in addition to WOC, HP1b, and HP1c, ROW binds to components of the insulator complex, BEAF-32, and Chromator. ChIP-seq experiments showed that ROW binds AT-rich sequences through three AT-hooks. The binding sites of ROW are located upstream to the transcription start sites of housekeeping genes and flanked by binding motifs of BEAF-32. Moreover, we found that the genome distribution of ROW was highly correlated with BEAF-32 and significantly enriched at TAD boundaries. Depleting *row* and *BEAF-*

*32* in S2 cells resulted in a correlated change in gene expression. The differential expressed genes were more likely to be downregulated, indirect targets of ROW and BEAF-32 (without binding sequences), and regulated through long-range contacts. The analysis of Hi-C data revealed enrichment of long-range interactions between promoters of housekeeping genes bound directly by ROW and promoters of developmental and inducible genes bound indirectly by ROW. Thus, our data show that ROW and BEAF-32 provide a general regulation mechanism depending on the contact between promoters of housekeeping and inducible genes.

## Results

### Knockdown of row causes a decrease in survival and fertility

To determine the functions of *row in vivo,* we utilized two publicly available UAS-*row*^RNAi^ transgenic fly lines, hereby referred to as *row*^RNAi-1^ and *row*^RNAi-2^. When combined with the ubiquitous *actin5C-GAL4* driver, the progenies carrying the Gal4 driver and *row*^RNAi^ construct show a significant decrease in ROW protein levels in fly heads, which was more substantial in *row*^RNAi-1^ relative to *row*^RNAi-2^ (94% and 87%, respectively; *P* <0.05; Fig. 1A).

**Figure 1.**
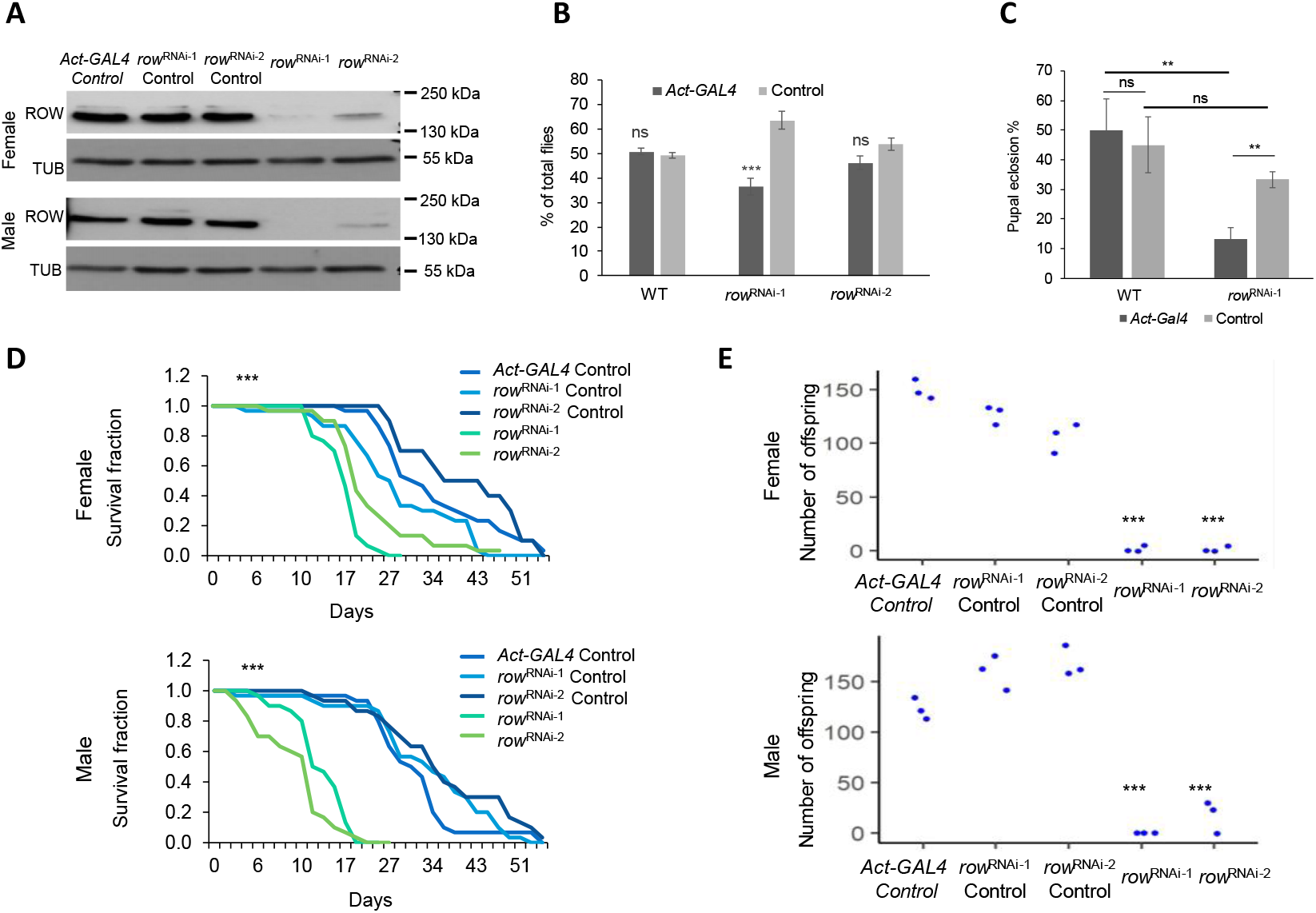
Effect of *row* knockdown on survival and fertility of *Drosophila melanogaster*. (A) Levels of ROW protein in *row*^RNAi^ lines were decreased in both male and female fly heads. Western blot for the two *row*^RNAi^ lines compared to controls was performed using rat polyclonal αROW and rat αTUBULIN (TUB). Genotype description: *row*^RNAi-1^(*Act-GAL4*/+; UAS-*row*^RNAi-1^/+), *row*^RNAi-2^(*Act-GAL4*/+; UAS-*row*^RNAi-2^/+), *row*^RNAi-1^ Control (Cyo/+; UAS-*row*^RNAi-1^/+), *row*^RNAi-2^ Control (Cyo/+; UAS- *row*^RNAi-2^) and *Act-GAL4* Control (*Act-GAL4* /+). (B) Decrease in the viability of *row*^RNAi^ lines. To examine the viability of *row*^RNAi^ flies, we counted the offspring generated from the cross between the heterozygous *Act-GAL4*/*Cyo* driver line with the homozygotes *row*^RNAi,^ or WT (w^1118^) flies as control. Values are the percentages from the total progeny ± standard error of the mean (SEM). (C) Decrease in pupal eclosion of *row*^RNAi^ line. Pupae were collected from the cross between the heterozygous *Act-GAL4*/*Cyo* driver line with the homozygotes *row*^RNAi-1^ or wild-type flies as control. Values are the percentages for each genotype of pupal eclosion ± SEM. (D) Survival curve for *row*^RNAi^ flies showing reduced life span compared to controls. (E) Crossing of females/males *row*^RNAi^ flies with WT males/females (w^1118^) flies resulted in a strong reduction in offspring number. Values are the number of offspring that resulted from crosses between WT flies and flies with different genotypes. Significance is represented by: **, *P* < 0.01; ***, *P* < 0.001; ns, non-significant. In D-E the significance level was received in all comparisons between each of the *row*^RNAi^ lines and the different controls.

We examined the viability of *row*^RNAi^ flies (Fig. 1B). Relative to the expected proportion of 50% (the Gal4 driver line is heterozygous for the insertion), there was a small but significant reduction in the progeny carrying both the Gal4 driver and expressing *row*^RNAi-1^, (36.4%; *P* < 1.1 ×10 ^-4^), but the reduction in viability was not significant for *row*^RNAi-2^ (46.2%; *P* =0.15; Fig. 1B). We observed that the lethality occurred at the pupal stage as *row*^RNAi-1^ pupal eclosion was significantly reduced (13% compared to 33% of *row*^RNAi-1^ control, *P*= 0.0058; Fig. 1C). In addition to development, knockdown of *row* also provoked shorted life span of the flies reaching adulthood for both *row*^RNAi^ lines (*P* < 0.001; Fig. 1D). In males, the effect on life span was very pronounced relative to the controls, while in females, we observed a smaller yet still significant life span reduction (Fig. 1D). To test the fertility of the flies lacking *row*, we collected virgin *males/females row*^RNAi^ flies, crossed them with wild-type (WT) females/males flies, and counted the number of offspring (Fig. 1E). Females and males *row*^RNAi^ flies showed a significant reduction in offspring number compared to control flies (*P* < 0.001 for both *row*^RNAi^). These results demonstrate that *row* expression is required for normal lifespan and reproduction, in addition to the previously described importance during development.

### Genes differentially expressed in row knockdown flies are associated with metabolism

To better understand the mechanisms responsible for the low fitness of the flies due to *row* knockdown, we generated and sequenced RNA-seq libraries from control and *row* knockdown fly heads (3-5 days old). As expected, the expression levels of *row* mRNA were significantly decreased in the *row*^RNAi^ lines by 40-60% (*P* < 0.01 for both *row*^RNAi^; Fig. S1A). The differential expression analysis showed that the effect of *row* knockdown on gene expression was consistent between the RNAi lines (r = 0.79, *P* < 2.2 ×10 ^-16^; Fig. 2A). However, the change in expression was more substantial for the line showing a more substantial *row* knockdown (*row*^RNAi-1^, the effect was on average 1.5 times larger; Fig. S1B-C). When analyzing the two *row*^RNAi^ lines together, we found 2035 genes with significant differential expression relative to the control (False Discovery Rate (FDR) < 0.05; Table S1). Despite the suspected role of ROW in transcriptional activations, the number of genes upregulated and downregulated was nearly equal (53% and 51% were upregulated in *row*^RNAi-1^ and *row*^RNAi-2^, respectively) (Fig. 2B and Fig. S1B-C).

**Figure 2.**
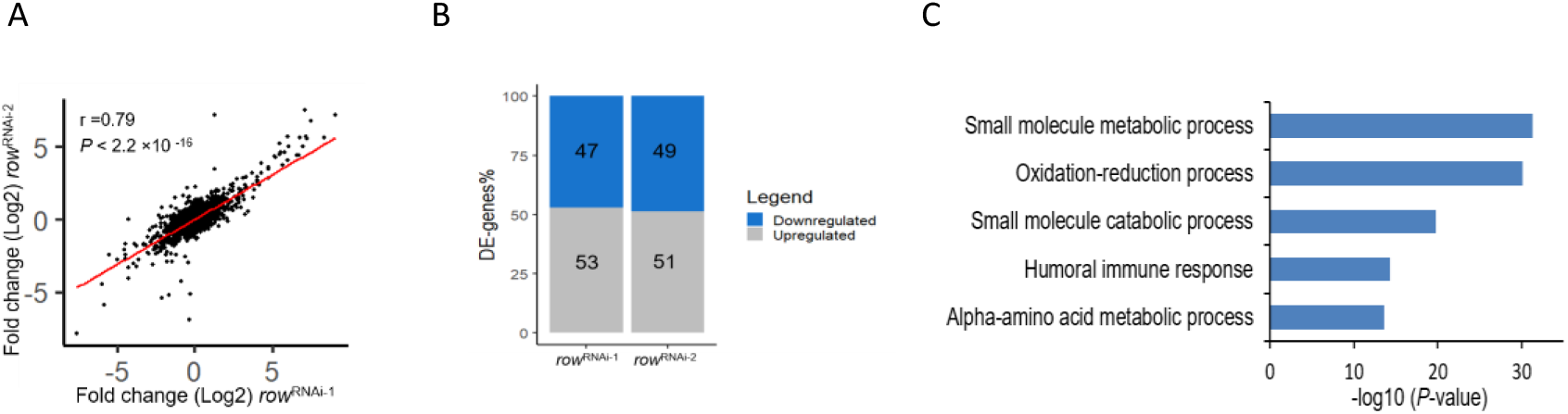
Knockdown of *row* in fly heads result in misregulation of genes involved in metabolism. (A) Correlation between the fold changes (log2) in *row*^RNAi-1^ and *row*^RNAi-2^. Fold changes were calculated relative to *Act-GAL4* control (using edgeR). (B) The percentage of differentially expressed genes (DE-genes) upregulated and downregulated in the two *row*^RNAi^ lines. (C) The top 5 most significantly enriched biological processes for differentially expressed genes after removing redundant terms.

We tested for enrichment of biological processes to examine what type of genes are differentially expressed in the *row*^RNAi^ lines. The most significant processes were related to metabolism, including oxidation-reduction processes and amino acid metabolism (Fig. 2C). This enrichment suggests that the fitness of the flies is reduced because of abnormal metabolism.

### ROW binds in vivo to HP1b/HP1c proteins and components of insulator complexes, BEAF-32 and Chromator

Our findings suggest a link between the chromatin protein ROW and the regulation of metabolism in vivo. To gain insights into the molecular mechanisms for the expression changes, we first studied the interactors of ROW in fly heads. We utilized CRISPR technology to FLAG-tag the endogenous ROW protein in flies. We validated the tagging with western blot (Fig. 3A) and sequencing (Fig. S2A). The tagging does not affect the levels of ROW protein (Fig. S2B). We then utilized these flies to identify ROW interacting proteins by performing affinity-purification of ROW-containing protein complexes from fly heads followed by mass spectrometry analysis. We found five co-purified proteins with ROW in all three independent experiments that were not detected in the control experiments (Fig. 3B). Another 32 proteins were co-purified with ROW in two out of the three experiments (Fig. S2C). The five most significant proteins were HP1c, HP1b, the zinc-finger finger protein WOC, the extraproteasomal ubiquitin receptor Dsk2 (also known as Ubqn), and the subunit of the cytoplasmic Dynein, Ctp (Fig. 3B). We used the molecular interaction search tool (MIST) (29) to identify protein interactions supported by additional evidence from previous studies. The analysis provided evidence for two highly connected complexes (Fig. S2D). The main complex that included ROW was composed of the five proteins that we identified as high-significant interactors (HP1c, HP1b, WOC, Dsk2, and Ctp), together with the transcription regulator *hfp*, and two components of an insulator complex: the boundary element BEAF-32 and the chromodomain protein, Chromator (Fig. 3B-C). To further confirm the interaction between ROW and BEAF-32, we performed Co-immunoprecipitation using S2 cells transfected with a ROW-FLAG tagged plasmid. Indeed, we found that immunoprecipitation of ROW using α-FLAG antibody results with coprecipitation of BEAF-32 (Fig 3D).

**Figure 3.**
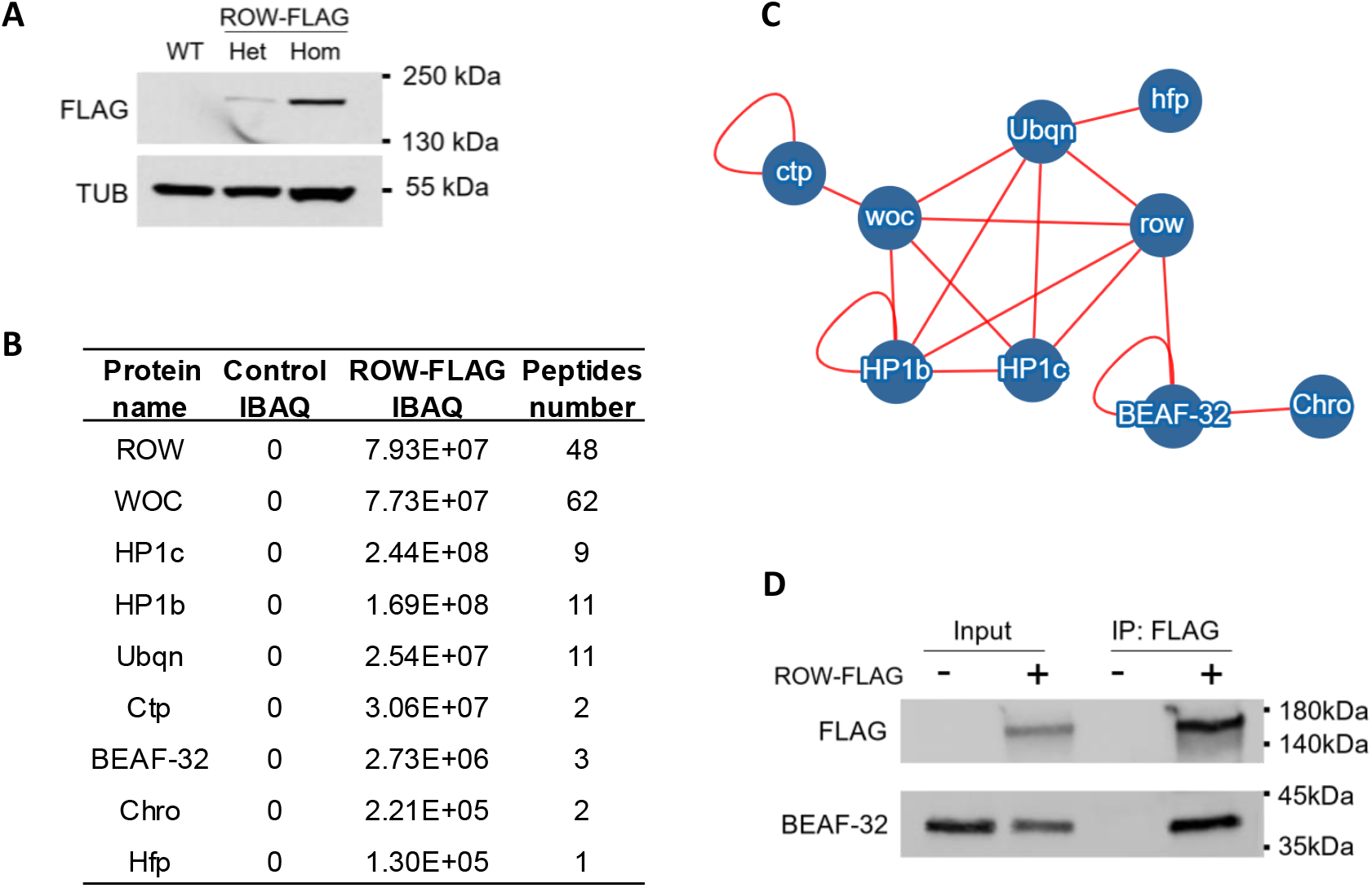
Characterization of the *in vivo* interactome of ROW in fly heads. (A) Western blot for flies expressing endogenous FLAG-tagged ROW (ROW-FLAG), with αFlag tag antibody to validate the tagging. αTubulin (TUB) antibody was used as a reference. Het, Heterozygotes; Hom, Homozygotes for tagged ROW. (B) Table summarizing the affinity purification–mass spectrometry data. IBAQ (intensity-based absolute quantification) reflects the protein abundance in the sample. Peptides number is the number of razor and unique peptides. The data are the mean for the control (W^1118^, n=3) and ROW-FLAG flies (n=3) samples. The proteins presented in the table were co-purified with ROW in at least two out of the three experiments, none in the control experiments, and are supported by additional evidence from previous studies. (C) Network of identified protein-protein interactions associated with ROW in fly heads. The red lines represent previously identified interactions. (D) BEAF-32 coimmunoprecipitate with ROW. Lysates from S2 cells not transfected (-) or transfected (+) with ROW-FLAG tagged plasmid were subjected to immunoprecipitation with αFlag tag beads. Immunoprecipitates and input (5%) were analyzed by western blot with αFlag tag and αBEAF-32 antibodies.

### ROW binds upstream to the transcription start sites of housekeeping genes but is less likely to bind genes that are differentially expressed by row knockdown

We assumed that the complex that includes ROW, WOC, HP1c, and HP1b is expected to be responsible for the transcription dysregulation in the fly head upon *row* knockdown. To test it, we performed ChIP-seq for ROW, WOC, HP1c, and HP1b in fly heads to identify the direct targets of the proteins in this complex. We initially examined the genome distribution relationship between the different proteins in the complex by calculating the pairwise correlation in ChIP-seq signals in non-overlapping bins across the genome. We found a very strong and significant correlation between the genome distribution of all the proteins in the complex (P < 2.2×10^-16^; Fig. 4A), but the most significant correlation was between ROW and WOC (r = 0.95) and between HP1c and HP1b (r = 0.94). We also examined the overlap between the binding sites that were identified for each protein (MACS2 peak caller, *q-value* < 0.05; the number of peaks: ROW = 5302, WOC = 4896, HP1c = 4252, HP1b = 2508). Similar to the quantitative analysis, we identified the strongest overlap of binding sites between ROW and WOC (82%) and between HP1c and HP1b (76%) (Fig. 4B). These results indicate that the core proteins of the ROW complex co-localize *in vivo* at an overlapping set of genomic positions. It also suggests that they may operate at specific binding sites as heterodimers formed by the assembly of ROW/WOC and HP1c/HP1b. We next characterized the location of ROW binding sites and found that most (69.8%) overlap promoter regions (*P*=9.9×10^-6^; Fig. 4C). We observed a slightly lower overlap with promoter regions for WOC (63.8%) and a substantially lower overlap for HP1c and HP1b (43.0% and 35.1%, respectively; Fig. 4C). The binding profile of all four proteins showed similar enrichment of approximately 150 bases upstream of the TSS (Fig. 4D). The ChIP-seq profile of the four proteins for a representative region is shown in Fig. 4E.

**Figure 4.**
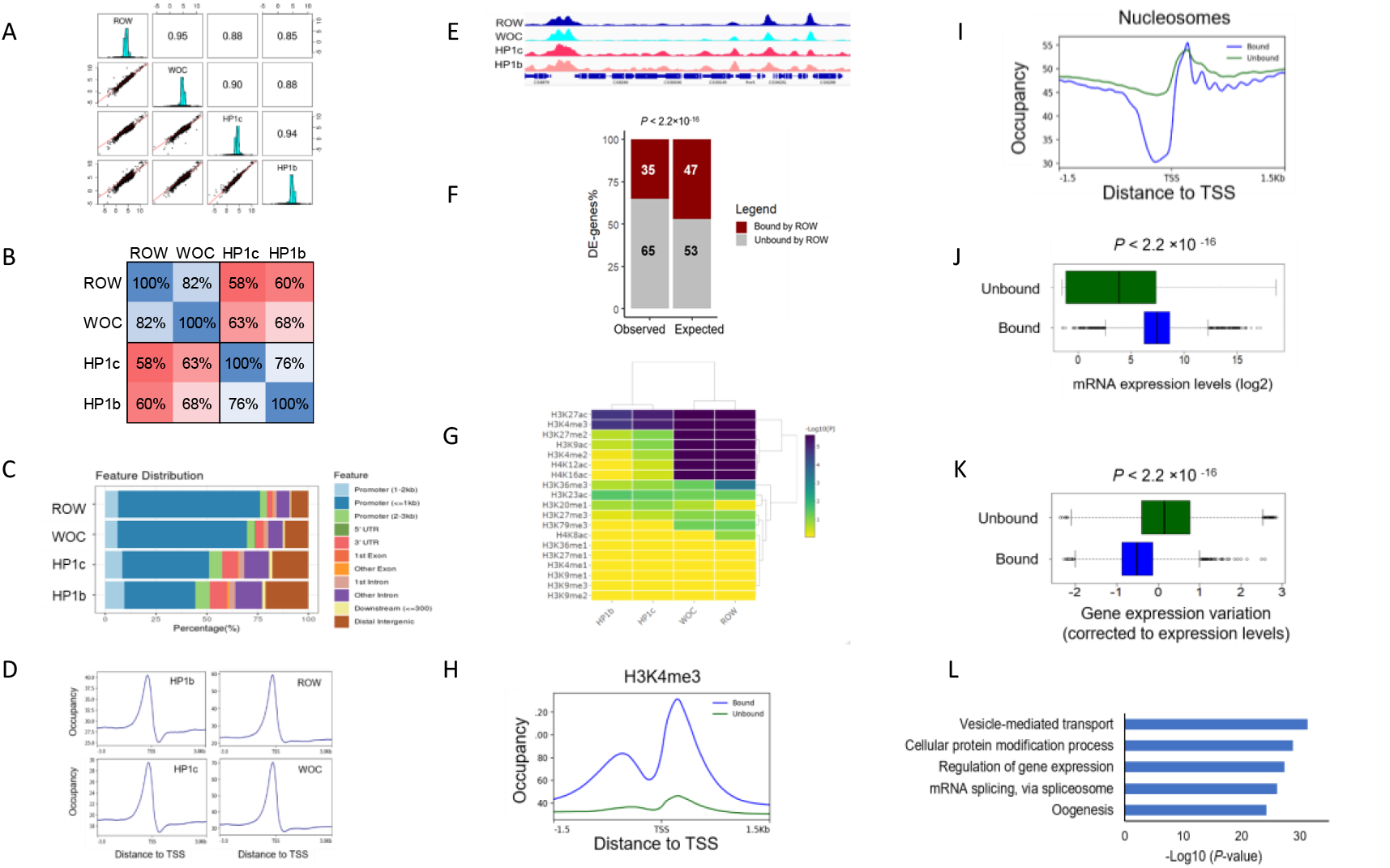
ROW binds *in vivo* to promoters of housekeeping genes. (A) Pairwise correlation between the ChIP-seq signals of ROW, WOC, HP1c, and HP1b in non- overlapping bins of 2000 bases across the genome. In the middle: the distributions of the ChIP- seq signal. On the bottom: bivariate scatter plots with linear regression lines are displayed. On the top: the value of the correlation. (B) Percentage of overlap between the binding sites of ROW, WOC, HP1c, and HP1b. (C) Genomic annotation of ROW, WOC, HP1c, and HP1b binding sites. (D) Average signal profiles (metagene plot) of ROW, WOC, HP1c, and HP1b over a 6-kb window around TSSs. (E) ChIP-seq signals for ROW, WOC, HP1c, and HP1b at example genomic region. (F) The percentages of differentially expressed genes in fly heads that were bound or unbound by ROW in fly heads and the expected values under an independence assumption. (G) Heatmap showing the significance of the overlap between different histone modifications and ROW, WOC, HP1c, and HP1b. (H) A metagene plot of H3K4me3 over a 3-kb window around TSSs for genes bound and unbound by ROW. (I) Average nucleosome profile over a 3-kb window around TSSs for genes bound and unbound by ROW. (J(Expression levels in the fly head for all genes bound and unbound by ROW. Values are log2 of the normalized reads count based on RNA-seq from control flies. (K) Gene expression variation (corrected to expression levels, see methods) for genes bound and unbound by ROW across 30 different developmental stages. (L) Enrichment of biological process for genes bound by ROW.

As ROW binds upstream to the TSS, we determined which genes are bound by ROW and found 4784 such genes (with ROW peak between - 250bp and +50 of the TSS) (Table S2). We then examined which of the differentially expressed genes in *row*^RNAi^ fly heads are bound by ROW. Surprisingly, out of 2035 differentially expressed genes, only 713 genes (35%) were bound by ROW, which is significantly lower than what is expected by chance (959 are expected by chance, P < 2.2×10^-16^; Fig. 4F). Thus, our analysis suggests that most differentially expressed genes upon row knockdown may represent indirect effects of the knockdown of *row*.

Given that the differentially expressed genes are mostly not ROW targets, we next asked what type of promoters and genes are bound by ROW. First, we tested the overlap between the ChIP-seq signal of the four proteins and histone marks (using modENCODE data (30)). We found that the histone marks of the active transcription, H3K4me3, and H3K27ac, showed a very significant overlap with all the four proteins in the complex (Fig. 4G). However, other histone marks that have been linked to active promoters (H3K4me2, H3K9ac, H4K16ac, and H4K12ac) showed very significant overlap only with ROW and WOC (Fig. 4G). To validate the results in fly heads, we performed ChIP-seq using an anti-H3K4me3 antibody. We compared the distribution of H3K4me3 near the TSS for genes bound by ROW relative to genes unbound by ROW and found a strong H3K4me3 signal flanking the peak of ROW (Fig. 4H).

Second, to further characterize the promoters bound by ROW, we performed micrococcal nuclease followed by sequencing (MNase-seq). The nucleosome structure can be used to distinguish between promoters of constitutively expressed genes (housekeeping genes) that typically show dispersed transcription from multiple TSSs in a relatively broad region (50-100 bases) with periodic arrays of nucleosomes and promoters of regulated genes with focused transcription, and lack organized chromatin structure (31–33). We found that the nucleosome organization of promoters bound by ROW was typical for active, housekeeping genes, with a wide nucleosome-free region (NFR) near the TSS and a regular array of nucleosomes upstream to the TSS (Fig. 4I).

Third, to establish that the genes bound by ROW are housekeeping genes, we examined their expression patterns and gene ontologies. Based on the RNA-seq we performed from fly heads, we found that ROW-associated genes show a significantly higher expression level than not-associated genes (*P* < 2.2 ×10 ^-16^; Fig. 4J). To test if ROW-associated genes are constitutively expressed, we studied expression variation across developmental stages and discovered that ROW-associated genes show low variation across stages (*P* < 2.2 ×10 ^-16^; Fig. 4K). Gene ontology (GO) analysis found that genes bound by ROW are enriched for multiple essential terms, including regulation of vesicle-mediated transport, regulation of gene expression, protein modification, mRNA splicing, and oogenesis (Fig. 4L).

In summary, our analysis indicates that ROW binds promoters of constitutively active genes, but those are less likely to be the genes that are differentially expressed by *row* knockdown.

### ROW binds AT-rich sequences through its AT-hook domains

To identify the DNA sequences responsible for ROW binding, we searched for enriched DNA motifs within the ROW binding sites. The most significant enrichment was AT-rich sequences (MEME-ChIP analysis: *E*-value = 3×10^-254^), located at the center of the ROW peak summit (Fig. 5A and Fig. S3A). We found a similar enrichment of AT-rich sequences for all the other proteins in the complex (WOC, HP1c, and HP1b; Fig. S3A). We used an available tool for predicting DNA-binding specificities for Cys2His2 zinc fingers (34) of ROW; however, the predicted sequences (Fig. S3B) did not resemble any motif that was significantly enriched in the binding sites of ROW. We thus predicted that the binding of ROW to AT-rich sequences is not through the zinc finger domains but more likely mediated through its AT-hook domains. To test this assumption, we performed ChIP-qPCR using αFlag tag antibody in S2 cells transfected with WT or AT-hook mutant ROW-FLAG tagged plasmids. The mutated version had a single amino-acid substitution in each of the 3 AT-hook domains of ROW (R105A, R632A, and R652A; Fig 5B). The WT and AT-hook mutant ROW expression in the transfected cells was similarly based on western blot (Fig S3C). Cell not transfected with ROW-FLAG tagged plasmid were used as a control. While ChIP-qPCR using cells transfected with WT ROW plasmid showed significant enrichment at the three promoters containing an AT-rich motif, the AT-hook mutant showed no significant enrichment at those promoters (Fig 5C). The results indicate that ROW binds to DNA by its AT-hook domains.

**Figure 5.**
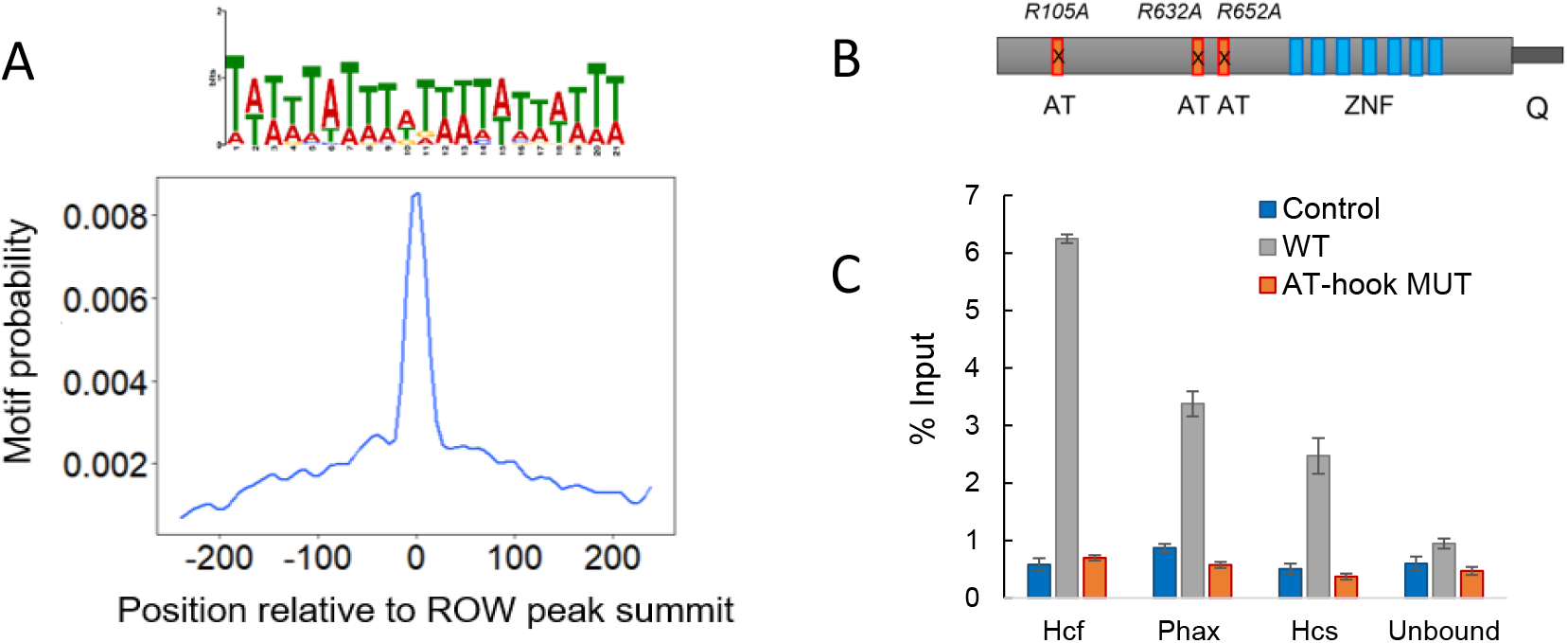
ROW binds specifically to AT-rich sequences by AT-hooks. (A) Central enrichment of AT-rich sequence in ROW binding regions. The logo shows the most significant enriched sequence (based on MEME-ChIP analysis). The plot shows the motif’s probability relative to the ROW ChIP-seq peaks (calculated with centriMo). (B) The structure of ROW is presented, with AT-hook domains (red), zinc-fingers (blue), and C-terminal glutamine (Q)-rich domain. The positions of the mutations performed in the AT-hook domains of ROW are shown. (C) ChIP-qPCR results using FLAG-tag antibody for S2 cells transfected with WT or AT-hook mutant ROW-FLAG tagged plasmids. As a control, cells not transfected with ROW-FLAG tagged plasmid were used. Input percentages are shown for three promoters containing AT-rich motif (Hcf, Phax, and Hcs) and a region unbound by ROW.

### ROW sites are flanked by binding sites and motifs of BEAF-32 at TAD boundaries

In addition to the AT-rich motifs, we found significant enrichment for a sequence motif (TATCGA) approximately 100bp from the peak summit of ROW (*E*-value = 8.4×10^-21^; Fig. 6A). Interestingly, BEAF-32 is known to bind this motif. To test whether ROW and BEAF-32 share binding sites across the genome, we compared the list of genes bound by ROW with those bound by BEAF-32 (identified previously in Kc167 cells (35) (Table S2). Indeed, 76.6% of the ROW-associated genes are also bound by BEAF-32 (Fig. 6B). The association between BEAF-32 and ROW is not only strong but highly specific, as an unbiased search for overlap between ROW-associated genes and genes targets of 84 transcription factors (36, 37) found that the most significantly enriched factor was BEAF-32 (Combined Score = 1255.3, FDR = 0) (Table S3). DREF binding motif is very similar to BEAF-32, but the overlap between DREF and ROW binding was not significant (Combined Score = 0.62, FDR = 0.068). These results suggest a substantial overlap between the binding of BEAF-32 and ROW. However, a metagene plot showed that BEAF-32 binding displayed a more pronounced peak closer to the TSS, suggesting that the two proteins act in proximity but not precisely in the same DNA location (Fig. 6C).

**Figure 6.**
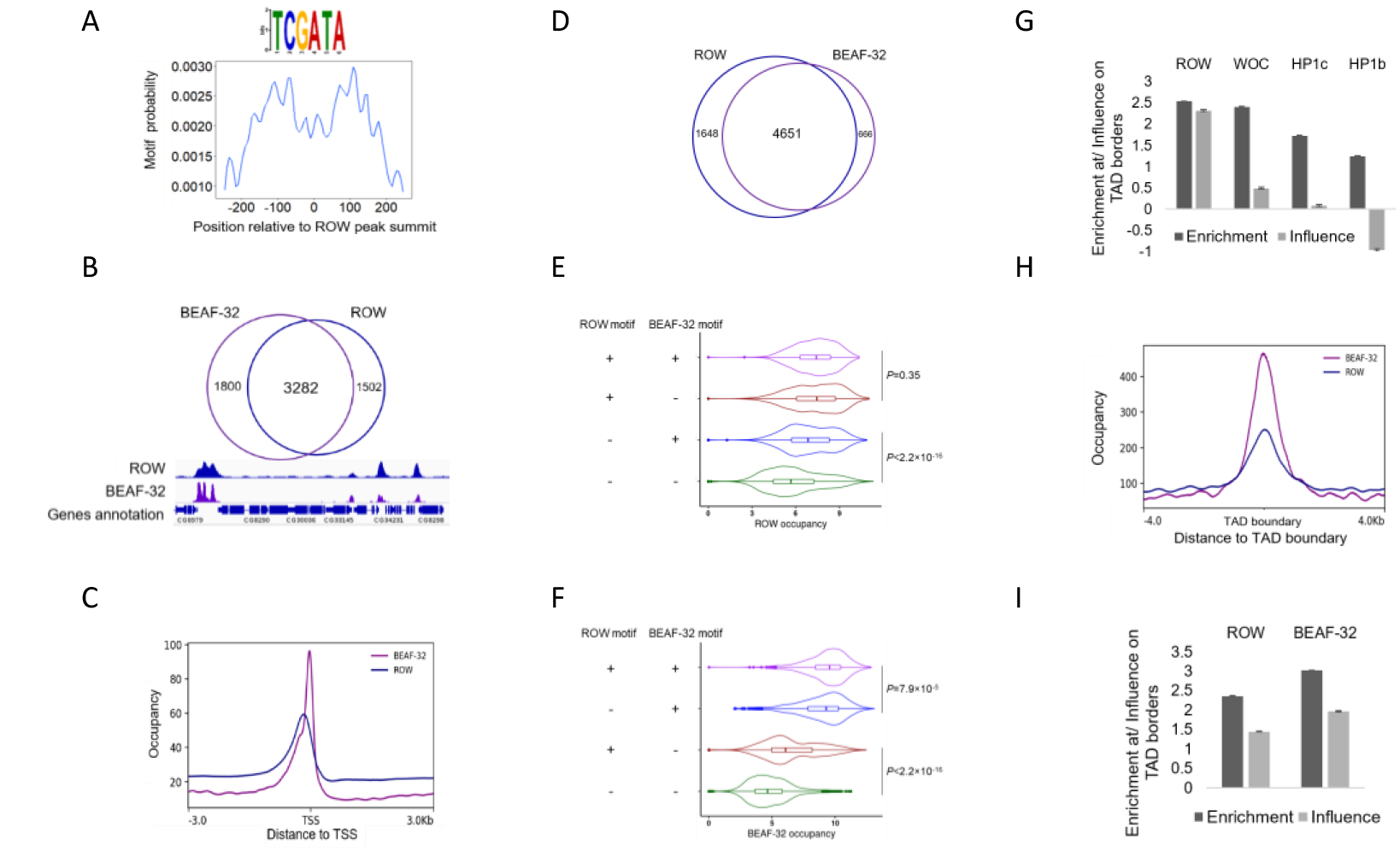
ROW and BEAF-32 ChIP-seq profiles relative to TSSs and TADs. (A) BEAF-32 consensus motifs (logo) were found to be enriched within ROW binding sites (MEME-ChIP analysis). The plot (calculated with centriMo) shows that the probability of having the BEAF-32 motif is highest around 100 bases from the center of the ROW ChIP-seq peaks. (B) Top: overlap between genes bound by ROW and genes bound by BEAF-32. Bottom: an example of ChIP-seq signal of ROW and BEAF-32 at a representative region with annotation of genes. (C) The distribution of ROW and BEAF-32 binding relative to the positions of TSSs. (D) The overlap between genes bound by ROW and genes bound by BEAF-32 in S2 cells. (E-F) The signal intensity of (E) ROW and (F) BEAF-32 at promoters with (+) or without (-) the binding motifs of ROW and BEAF-32 (based on ChIP-seq results in S2 cells). (G) Multiple logistic regression was used to compare the enrichment and independent influence of the proteins (ROW, WOC, HP1c, and HP1b) at TAD boundaries. Values are the enrichment and influence beta coefficients ± standard error calculated by the HiCfeat R package (38). (H) The distribution of ROW and BEAF-32 binding relative to the positions of TAD boundaries (based on ChIP-seq results in S2 cells). (I) Results of multiple logistic regression that was used to compare the enrichment and independent influence at TAD boundaries between BEAF-32 and ROW (based on ChIP-seq results in S2 cells). Values are the enrichment and influence beta coefficients ± standard error (calculated by HiCfeat R package (38)).

To compare the genome distribution of ROW and BEAF-32 in the same cells, we performed ChIP-seq for ROW and BEAF-32 in S2 cells. We replicated in this experiment the enrichment of AT-rich sequences and BEAF-32 consensus motifs in the binding sites of both proteins (Fig. S4A-B). We also observed an even more substantial overlap between the genes bound by the two proteins (87.5% of genes bound by BEAF-32 were bound by ROW and 73.8% vice versa) (Fig. 6D and Table S4). Similar to the *in vivo* data, the ChIP-seq signals of ROW and BEAF-32 were slightly shifted relative to TSSs (Fig. S4C and E).

Our findings show that ROW and BEAF-32 interact physically and bind the same promoters, but each protein binds a different sequence. We, therefore, asked if the ChIP-seq signals are consistent with the possibility that ROW and BEAF-32 assist in the recruitment of each other to specific sites. To test it, we checked how the binding intensity of ROW and BEAF-32 at gene promoters is influenced by ROW and BEAF-32 motifs. We found that for sites without the ROW AT-rich sequence (15 repeats of A or T), the binding intensity of ROW is significantly higher at promoters with BEAF-32 motifs (TCGATA) compared to promoters without it (*P* < 2.2 ×10 ^-16^; Fig. 6E). A similar result was observed for BEAF-32. For sites without BEAF-32 motifs, the binding intensity of BEAF-32 was higher at promoters with ROW motif than promoters without it (*P* < 2.2 ×10 ^-16^; Fig. 6F). Moreover, the binding intensity of BEAF-32 at promoters with ROW and BEAF-32 motifs is higher than promoters with only BEAF-32 motifs (*P* =7.9 ×10 ^-5^; Fig. 6F). These results suggest that BEAF-32 and ROW may assist in recruiting each other to specific promoters.

Since BEAF-32 occupies TAD boundaries in *Drosophila* (4), we tested the association of ROW (based on the ChIP-seq in fly heads) with TAD boundaries (7). Indeed, we found that ROW is enriched at the boundaries of TADs (enrichment coefficient *β̂* = 2.54, *p* < 1×10^−20^; Fig. 6G). The other partners of the ROW complex showed enrichment at TAD boundaries at a lower level (*β̂* = 2.41, 1.73, 1.24 for WOC, HP1c, HP1b, respectively; Fig. 6G). Since the proteins in the complex show correlation in their genome distribution (Fig. 4A-B), and this correlation may explain the enrichment at TAD boundaries (Fig. S4D), we used multiple logistic regression to test for the independent influence of each protein, as previously performed (38). We found a large decrease in the beta coefficients (estimates the independent influence of the variable on the outcome) for all the proteins except ROW (Fig. 6G). These findings suggest a specific and independent role for ROW at TAD boundaries and that the enrichment of WOC, HP1c, and HP1b at TAD boundaries is due to their correlation with ROW.

To compare the binding of ROW and BEAF-32 relative to TAD boundaries, we used our ChIP-seq in S2 cells. The signals of ROW and BEAF-32 were both centered on TAD boundaries (Fig. 6H and Fig. S4E). The enrichment at TAD boundaries was higher for BEAF-32 (*β̂* = 3.02, *p* < 1×10^−20^) relative to ROW (*β̂* = 2.36 *p* < 1×10^−20^; Fig. 6I). Multiple logistic regression showed a proportional reduction of the beta enrichment coefficient for both BEAF-32 (*β̂* = 1.97, *p* < 1×10^−20^) and ROW (*β̂* = 1.45, *p* < 1×10^−20^; Fig. 6I). This indicates a similar independent enrichment of BEAF-32 and ROW at TAD boundaries and suggests that the combination of ROW and BEAF-32 signals can better predict TAD boundaries.

### ROW and BEAF-32 regulate the expression of genes that are indirect targets

Since ROW and BEAF-32 bind most of the same promoters, we wanted to examine if they also have similar effects on transcription. We treated the S2 cells with dsRNA against *row*, *BEAF-32,* or both genes and analyzed gene expression in treated and untreated cells with RNA-seq. The reduction in protein levels was verified using western blot (Fig. 7A). We found a significant correlation between the changes in expression in the cells with *BEAF-32* knockdown and *row* knockdown (r = 0.44, *P* < 2.2×10^-16^; Fig. 7B), suggesting an overlap in the genes influenced by ROW and BEAF-32. The effect on gene expression was the strongest when both genes were knockdown, and it was the weakest in cells with only BEAF-32 knockdown (*P* = 0.017; Fig S5A). Changes in gene expression in cells with knockdown of both *row* and *BEAF- 32* were significantly correlated with the changes in cells with knockdown of *row*, knockdown of *BEAF-32*, and with the additive effect of the two genes (the sum of the fold changes in *row* and *BEAF-32* separate knockdowns) (Fig. S5B).

**Figure 7.**
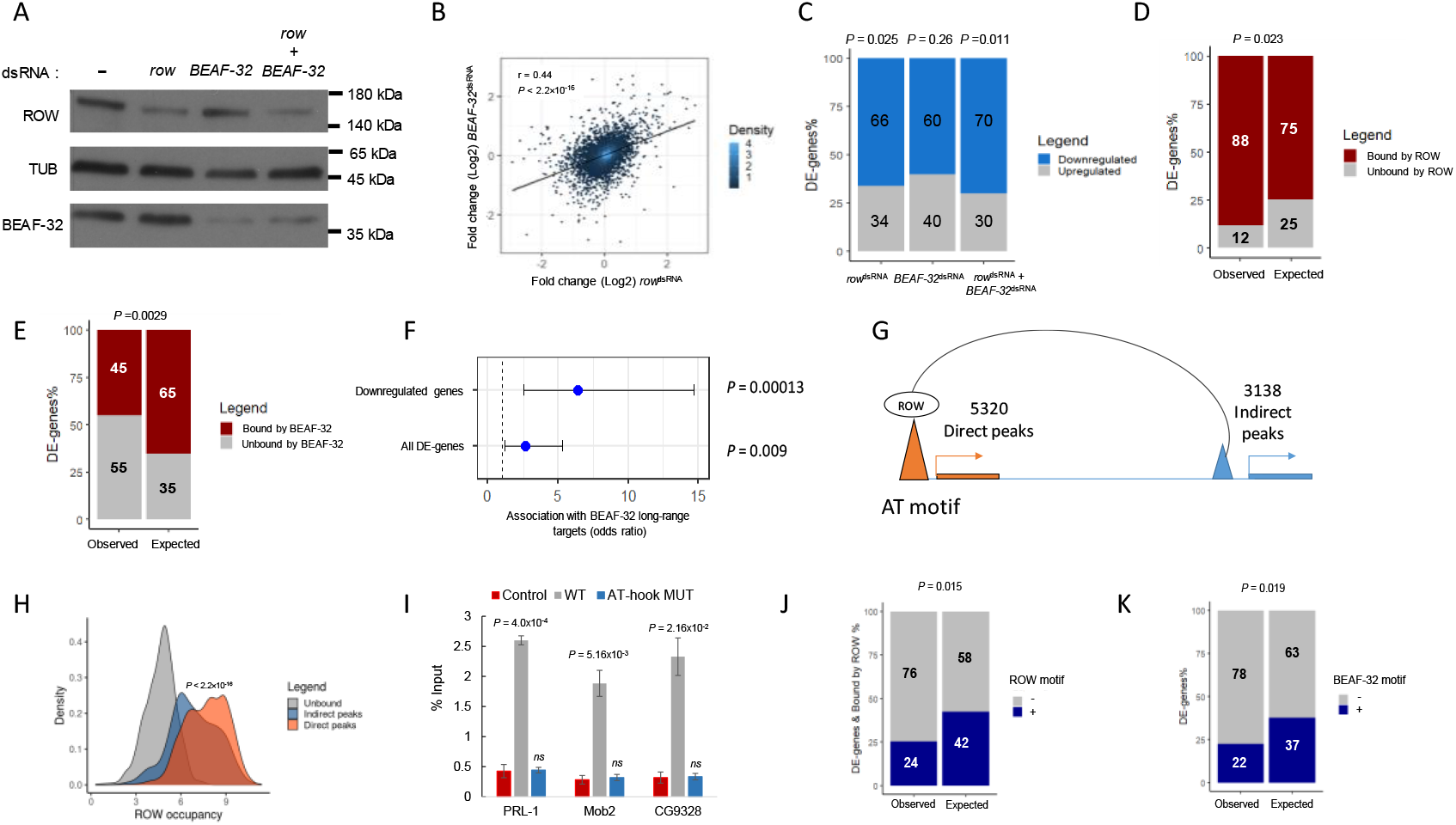
Knockdown of *row* or BEAF-32 in S2 cells causes downregulation of a common set of long-range targets. (A) Protein levels of ROW, BEAF-32, and Tubulin (TUB) in non-treated S2 cells and cells treated with dsRNA against *row*, *BEAF-32,* or both. _(B)_ Correlation between the gene expression fold change (log2) in cells treated with *row*^dsRNA^ and cells treated with *BEAF-32*^dsRNA^. The fold change was calculated relative to control cells. (C) The percentage of differentially expressed genes (DE-genes) upregulated and downregulated in *row*^dsRNA^, *BEAF-32*^dsRNA^, and *row*^dsRNA^ & *BEAF-32*^dsRNA^ treated cells. (D-E) The percentages of differentially expressed genes bound or unbound by ROW (D) or BEAF-32 (E) and the expected values under an independence assumption. (F) Association between the differentially expressed genes in S2 cells with *row* and BEAF-32 knockdown and previously published long-range targets of BEAF-32. Values are the odds ratio ± 95% confidence interval of the association between all genes or restricting the analysis to downregulated genes. (G) Illustration of ROW direct and indirect peaks at promoter regions. (H) Density plots of ROW binding signal (ChIP-seq) at promoters of genes unbound by ROW, promoters bound directly by ROW (with AT-rich motif), and promoters bound indirectly by ROW (without AT-rich motif). (I) ChIP-qPCR results using FLAG-tag antibody for S2 cells transfected with WT or AT-hook mutant ROW-FLAG tagged plasmids. As a control, cells not transfected with ROW-FLAG tagged plasmid were used. Input percentages are shown for three promoters with an indirect peak of ROW (PRL-1, Mob2, and CG9328). (J) The percentages of differentially expressed genes bound by ROW with or without a ROW motif (15 repeats of A or T) at the promoter region and the expected values under an independent assumption. (K) The percentages of differentially expressed genes with or without a BEAF-32 motif (TCGATA) at the promoter region and the expected values under an independence assumption.

Comparing gene expression in the treated and untreated cells, we found 51 differentially expressed genes (FDR < 0.1; Table S5). The 51 genes had a significant overlap with a previously reported list of differentially expressed genes upon *row* knockdown in S2 cells (22) (OR = 24.1; *P* = 3.2×10^-13^). The majority of the differential expressed genes in the S2 cells were downregulated (*row* knockdown: 66%, *P* = 0.025; *BEAF-32* knockdown: 60%, *P* = 0.26; knockdown of both: 70%, *P* = 0.011; Fig. 7C), consistent with the role of ROW and BEAF-32 in transcription activation. We used our ChIP-seq results to test if the differentially expressed genes are bound by ROW and BEAF-32. We found a positive association between the differentially expressed genes and binding by ROW (88% bound by ROW vs. 75% expected by chance, *P* = 0.023; Fig. 7D), but surprisingly for BEAF-32, the association was significantly lower than expected by chance (45% bound by BEAF-32 vs. 65% expected, P=0.0029; Fig. 7E).

The negative association between BEAF-32 binding and expression changes could result from the involvement of BEAF-32 in transcription regulation through long-range contacts between direct ChIP peaks of BEAF-32 (containing DNA binding motifs) and indirect low-intensity peaks (without BEAF-32 motifs), as was previously described (10). To test it, we compared the list of the 51 differentially expressed genes that we identified with a previously published list of genes that BEAF-32 regulates in S2 cells through long-range contacts (10). Those genes were found by introducing a mutation that impairs the interaction between BEAF-32 and CP190, which abolishes the binding of BEAF-32 to the indirect peaks (10). Remarkably, we found a significant association between the 51 differentially expressed genes and the long-range targets of BEAF-32 (OR = 2.6, P = 0.009; Fig. 7F), which was more significant for downregulated genes in both data sets (OR = 6.4, P = 0.00013; Fig. 7F). This finding implies that the downregulated genes we identified are activated through the long-range and indirect binding of BEAF-32.

As an interactor of BEAF-32, we speculated that ROW might also have indirect low-intensity peaks associated with changes in expression. These low-intensity peaks could have been missed in our *in vivo* ChIP-seq and could explain why the differentially expressed genes in the fly head were not associated with ROW binding. We, therefore, divided the peaks of ROW in the S2 cells ChIP-seq to direct peaks with AT-rich sequences (15 repeats of A or T) and indirect peaks without AT-rich sequences, which included 5320 direct peaks and 3138 indirect peaks (Fig 7G). The majority of both direct and indirect peaks of ROW overlap promoters (60.3% and 65.5%, respectively) (Fig. S5C). As predicted, the promoters with indirect peaks had a lower intensity than promoters with direct peaks (*P* < 2.2×10^-16^; Fig. 7H). To confirm that the indirect ROW peaks are specific, we performed ChIP-qPCR on three promoters with indirect peaks using cells transfected with WT ROW plasmid and the AT-hook mutant. The three sites were specifically enriched by immunoprecipitation with WT ROW, but not with the AT-hook mutant, showing that indirect peaks are a result of ROW binding to AT-rich sequences and probably through long-range contacts between direct and indirect peaks (Fig 7I).

Next, we tested if the differentially expressed genes in the S2 cells treated with dsRNA against *row* and *BEAF-32* are indirect ROW targets. Although most of the differentially expressed genes are bound by ROW, only 24% have the ROW motif (42% are expected by chance; *P* = 0.015; Fig. 7J). We also found that the differentially expressed genes are depleted in the BEAF-32 motif (22% have the motif vs. 37% expected by chance; *P* = 0.019; Fig. 7K). This suggests that the binding of ROW to the promoter of the differentially expressed genes is the outcome of indirect binding.

### ROW binds directly to housekeeping genes and indirectly to developmental genes via long-range interactions

Our analysis showed that ROW binds housekeeping genes, but it is not clear what type of genes are directly and indirectly bound by ROW. We analyzed the enrichment of GO terms for genes bound directly and indirectly by ROW to test it. We found that genes bound indirectly by ROW are enriched with GO terms related to developmental and regulated processes (Fig. 8A), while genes bound directly by ROW were enriched with processes involved in basic maintenance of the cells (i.e., housekeeping genes; Fig. 8B).

**Figure 8.**
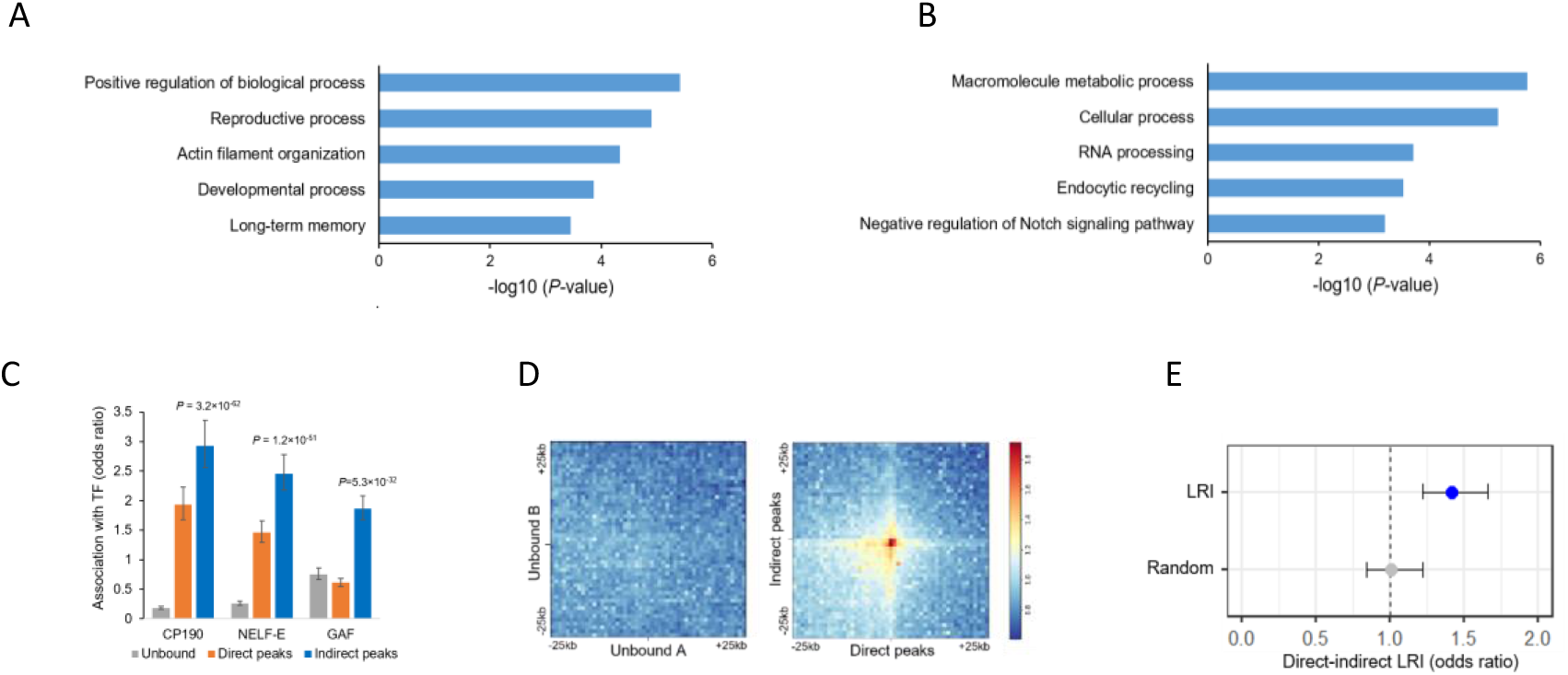
Long-range interactions between promoters of housekeeping genes directly bound by ROW and developmental genes bound indirectly by ROW. (A-B) Enrichment of biological process for (A) genes bound indirectly and (B) directly by ROW. (C) The association between genes unbound, bound directly, and bound indirectly by ROW with three transcription factors (TF): CP190, NELF-E, and GAF. Values are the odds ratios ± 95% confidence interval. (D) Plot of aggregated Hi-C sub-matrices of (Left panel) random sets of promoters unbound by ROW, and (Right panel) promoters bound directly and indirectly by ROW. The plots show the promoters in the center within a region of 50kb divided into 50 bins (bin size = 1kb). The values are the mean of observed/expected transformed sub-matrices (warm colors indicate higher values). The middle region on the right panel shows a high observed/expected value indicating a high level of long-range interactions between promoters bound directly and indirectly by ROW. (E) Significant enrichment of long-range interactions (LRI) between direct and indirect ROW peaks, but not between randomly generated interactions. The association tests within the interactions identified were performed in Hi-C data using the PSYCHIC tool (41). Values are the odds ratios ± 95% confidence interval.

Long-range targets of BEAF-32 were previously found to be enriched with factors associated with RNA pol II pausing (GAF and NELF) (10), and CP190 was found to be required for long-range interactions of BEAF-32. Promoters with highly paused RNA pol II tend to be of developmental genes often bound by GAF, which is essential for establishing paused Pol II (39, 40). Consistent with this observation, indirect peaks of ROW were positively associated with CP190 (OR = 2.9, *P* = 3.2×10^-62^), NELF-E (OR = 2.5, *P* = 1.2×10^-51^), and GAF (OR = 1.9, *P* = 5.3×10^-32^; Fig. 8C). Direct peaks of ROW were also enriched, to a lesser extent, with CP190 (OR = 1.9, *P* = 6.7×10^-22^) and NELF-E (OR = 1.5, *P*=1.1×10^-9^), but were negatively associated with GAF (OR = 0.6, *P* = 2.6×10^-17^; Fig. 8C).

To validate the occurrence of the long-range interactions between promoters bound directly and indirectly by ROW, we used genome-wide aggregation of published Hi-C data in S2 cells (7). While a random set of promoters unbound by ROW showed no enrichment for long-range interactions (mean observed/expected = 0.99; Fig. 8D, left), promoters bound directly and indirectly by ROW had high levels of long-range interactions (mean observed/expected = 1.8; Fig. 8D, right). To further confirm the existence of the long-range interactions, we used a computational approach that identifies over-represented promoter interactions in Hi-C data (41). There was a significant association between direct and indirect ROW peaks within these identified interactions (OR = 1.42, *P* = 6×10^-6^; Fig. 8E). Thus, our results demonstrate the existence of long-range interactions between promoters of housekeeping genes, bound directly by ROW, and promoters of developmental genes, bound indirectly by ROW.

## Discussion

Long-range chromatin interactions have an essential role in transcription regulation. Our data strongly indicate that *row* is required for the transcription regulation of developmental and inducible genes by forming promoter-promoter interactions with housekeeping genes. Our study uncovers new cooperation between the insulator protein BEAF-32 and the chromatin binding protein ROW (Figure 9). The two proteins interact and bind to the same genomic positions, which are promoters of housekeeping genes located at the boundaries of TADs. The ROW binding sites are AT-rich sequences flanked by motifs and binding of BEAF-32. The association of BEAF-32 and ROW with TAD boundaries may be an indirect effect of the localization of the two proteins at TSSs of active genes. While ROW directly binds promoters of housekeeping genes, we found that the knockdown of *row* affects the expression of genes that are long-range targets indirectly bound by ROW. The existence of long-range interactions between housekeeping genes bound directly by ROW and inducible genes bound indirectly by ROW appears to play an important role in gene regulation, making row an essential gene.

**Figure 9.**
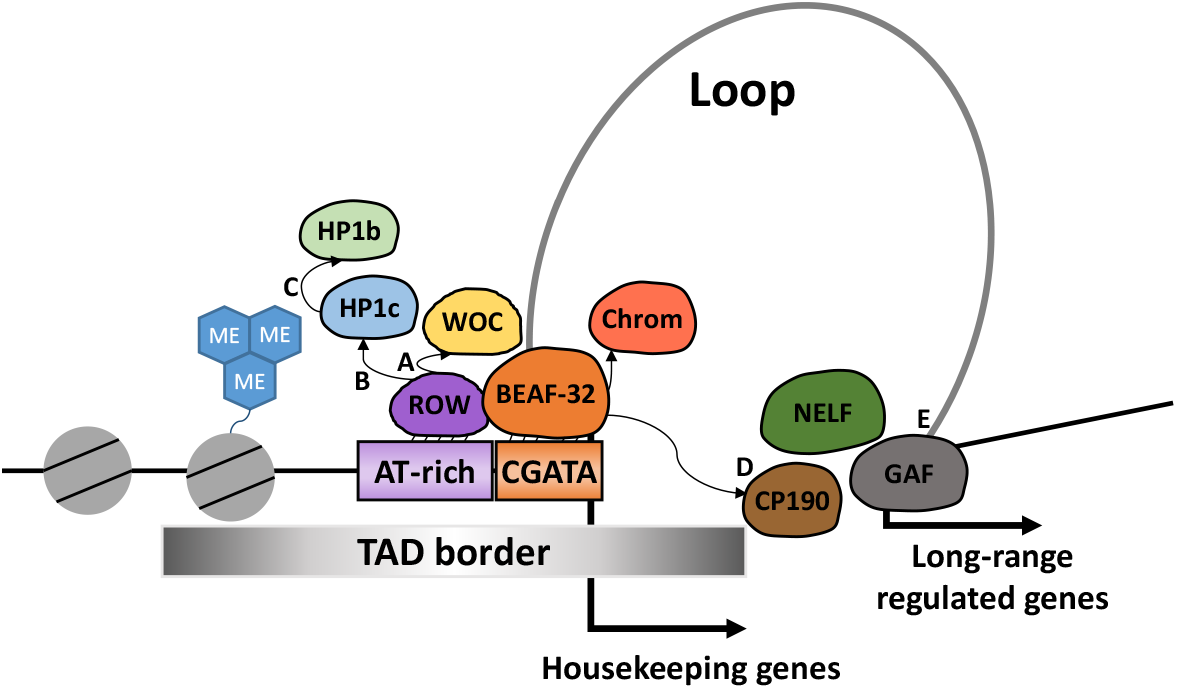
A model for the cooperation of ROW with BEAF-32 and other chromatin-bound proteins in transcription and nuclear organization. Our proposed model is that ROW and BEAF-32 provide DNA specificity for the two complexes. ROW recruits the HP1c complex previously shown to be involved in gene activation (22). BEAF- 32 interacts with CP190 and Chromator, responsible for the physical interactions required for long-range contacts (44). The model relies on the interaction between ROW and BEAF-32, the localization of both proteins at most of the same promoters, and the presence of AT-rich sequences and CGATA motifs in the promoters of housekeeping genes. Those promoters are characterized by a broad nucleosome-free region and high levels of H3K4me3. The recruitment of proteins such as WOC and HP1c to the site by ROW is proposed to be essential for transcription activation of developmental genes that have long-range contact with the active promoters. (A) Our study shows that WOC has the highest overlap in binding with ROW. A previous study showed that ROW is required for WOC binding to the chromatin (24). Our study shows that ROW binds selectively to AT-rich sequences using AT-hooks motifs. (B) HP1c shows a lower correlation with ROW relative to WOC. HP1c binding to the chromatin is also dependent on ROW (20, 24). (C) The correlation in genome distribution is highest for HP1b with HP1c. Among the proteins tested, HP1b shows the lowest correlation with ROW and with the genomic annotations associated with ROW (active promoters and TAD boundaries). A previous study showed that HP1c and HP1b form heterodimers *in vitro* and *in vivo* (43). (D) Previously published studies showed that BEAF-32 regulates the expression of genes not directly bound by BEAF-32 through long-range contacts that depend on the interaction of BEAF-32 with CP190 (10, 11). (E) Based on our study, ROW and BEAF-32 regulate a common set of genes that are indirect targets of the two proteins (promoters that lack the DNA binding motifs). We also show the occurrence of long-range contacts between promoters of housekeeping genes and developmental genes. GAF, NELF, and CP190 bind the long-range targets of ROW and BEAF- 32

Our results show that the hierarchical recruitment of the protein complex that includes HP1b/c and WOC to active promoters depends on the sequence-specific binding of ROW and the interaction with BEAF-32. BEAF-32 recruits other insulator proteins to regulate genes through long-range interactions (Figure 9). Based on our findings, the specificity of the binding of the multiple proteins involved in the long- range regulation is due to the cooperation of two sequence-specific DNA-binding proteins: BEAF-32 and ROW. BEAF-32 binds to its motifs (CGATA) (42), located near AT-rich sequences bound by ROW. The binding of ROW to AT-rich motifs is mediated by its AT-hook domains. Our analysis suggests that the binding of ROW and BEAF-32 is facilitated by each other. After ROW and BEAF-32 bind the DNA, they may recruit other proteins, like WOC and HP1c by ROW and CP190 or Chromator by BEAF-32. The notion that the sequence-specific binding of ROW and BEAF-32 directs the localization of the HP1c complex is consistent with the strongest enrichment of ROW to promoters (relative to WOC, HP1c, and HP1b), and a more significant independent enrichment of BEAF-32 and ROW to TAD boundaries. Previous studies showed that the recruitment of WOC and HP1c to the chromatin is dependent on ROW (20, 24), but our results show a much higher concordance in the binding of WOC and ROW and a much lower correlation between ROW and HP1c. The recruitment of HP1b may be dependent on the heterodimerization with HP1c, as evident by the high correlation we observed in the genomic distribution between HP1c and HP1b and the lowest correlation of HP1b with ROW. This is consistent with the findings that HP1c/HP1b heterodimers are formed both *in vitro* and *in vivo* (43).

It was previously found that the ROW complex binds developmental genes associated with RNA pol II pausing (22). We found that the binding of ROW to developmentally regulated genes is established indirectly through long-range contacts with direct binding sites at promoters of housekeeping genes. Our work defines the ROW protein complex as essential for transcriptional activation of developmental and inducible genes. The mechanism that may explain how the complex of ROW can promote transcriptional activation of long-range genes is through stabilizing the NELF complex and stalled Pol II at those promoters. The depletion of *Dsk2,* a binding partner of ROW, causes a decrease in NELF-E and Pol II pausing at TSSs (22). Stalled Pol II is known to enhance the expression of developmental genes by maintaining accessible chromatin structure (45, 46). In addition, the recruitment of the complex FACT can facilitate RNA Pol II elongation. A previous study showed that HP1c recruits FACT to active genes and active forms of RNA polymerase II, and in the absence of HP1c, the recruitment of FACT into heat-shock genes was altered, and the expression levels were reduced (25).

In conclusion, the above results demonstrate an essential role for ROW and the HP1b/c complex in transcription activation of developmental genes through long-range interactions with promoters of housekeeping genes and add a new dimension to our understanding of the relationship between genome organization and transcription.

## Materials and Methods

### Fly stocks and maintenance

The *row*^RNAi-1^ and *row*^RNAi-2^ are 25971 and v28196, respectively, from the Bloomington *Drosophila* Stock Center and VDRC Stock Center. *Act-GAL4*/CyO flies (3953) were ordered from the Bloomington Stock Center. All flies were raised at 25°C in 12-hour light: 12-hour dark cycles on standard diets. We generated an isogenic background for the two *Drosophila row*^RNAi^ strains by crossing females seven times with W^1118^ male flies. In each cross, female offspring carrying the transgene were selected by the red-eye phenotype or by genotyping.

### Antibodies

Polyclonal rat αROW (20) (WB 1:10000), polyclonal rabbit αROW (22) (ChIP-seq: 1µl), αHP1c (22) (ChIP-seq: 5µl) and αHP1b (22) (ChIP-seq: 5µl) were a gift from the lab of Prof. Fernando Azorin. Rabbit αWOC (47) (ChIP-seq: 1µl) was a gift from the lab of Prof. Maurizio Gatti. Mouse monoclonal αFlag is SIGMA ALDRICH (F1804, WB 1:500). Rat monoclonal αTubulin (WB: 1:10000) and rabbit αH3K4me3 (ChIP-seq: 5µl) are Abcam (ab6160 and ab8580 respectively). Rabbit igG is Santa Cruz (sc-2027, ChIP-seq: 5µl). Mouse αBEAF-32 is Developmental Studies Hybridoma Bank (AB_1553420, WB: 1:200, ChIP-seq: 8µl).

### Western blot

20 fly heads per sample were collected on dry ice and homogenized in 200µl RIPA lysis buffer (50 mM Tris-HCl at pH 7.4, 150 mM NaCl, 1 mM EDTA, 1% NP-40, 0.5% sodium deoxycholate, 0.1% sodium dodecyl sulfate, and 1 mM DTT, and protease inhibitor tablets [Roche]) by motorized pestle. For S2 cells, 10^6^ cells were homogenized in 100 µl of RIPA buffer. The lysates were kept on ice for 20 min and then centrifuged at max speed for 30 minutes. The supernatants were collected, and 20µl per sample was boiled with protein sample buffer (Bio-Rad). Criterion XT Bis-Tris gels (Bio-Rad) were used for gel electrophoresis.

### Viability assay

The UAS-*row*^RNAi^ flies or W^1118^ control flies were crossed with *Act-GAL4*/CyO flies. Then, the number of CyO and non-CyO offspring were counted. The percentage of the specific genotype from the total progeny was calculated. Each cross was performed three times, and the result’s significance was tested using a two-sided binomial test.

### Pupal eclosion

The UAS-*row*^RNAi-1^ flies or control W^1118^ flies were crossed with *Act-GAL4*/CyO flies (n=3 for each cross-type). Seven days after the cross was made, 40 pupae were randomly collected, each pupa to a separate tube with food and a hole for fresh air. The number and the genotype of newly enclosed flies were recorded. The Statistical test performed was a two-sided t-test.

### Survival assay

From each genotype, three vials containing 20 flies (0-3 days old, ten males and ten females) were kept on standard food at 25°C. Once in three days, the number of flies that died was counted, and their gender was recorded. The experiment was continued until all flies died. Tests for the difference between the survival curves were performed with the OIsurv R package using the survdiff function.

### Fertility assay

Eight males or 12 females were crossed with w^1118^ flies from each genotype, and the number of offspring was counted. Three crosses were performed for each genotype. Statistical tests were performed with ANOVA followed by Tukey’s test.

### Plasmid generation and transfections for Co-immunoprecipitation (Co-IP) and ChIP-qPCR in S2 cells

To overexpress ROW in S2 cells, we generated ROW FLAGx2-tagged plasmid controlled by P-MT (Metallothionein) promoter (pMT-ROWx2FLAG). pMT-V5 plasmid (Invitrogen) was cut using KpnI and XhoI. The gene ROW was amplified by PCR using the RE01954 (BDGP) plasmid as a template. The primers included two FLAG-TAG sequences. As we used the Gibson Assembly kit, the primers included part of the plasmid backbone as well.

The primers used for the cloning:

5’-AGGGGGGATCTAGATCGGGGTACAGTTAGCTGTAAGATGACGC-3’ (F)

5’-CTTCGAAGGGCCCTCTAGACTCACTTGTCATCGTCATCCTTGTAATCCTTGTCA

TCGTCATCCTTGTAATCCAATTGCGGATGGTGATGGTG-3’ (R)

For generating ROW FLAGx2-tagged plasmid with mutations in the AT-hook domains, PCR was performed using the WT plasmid and primers that contain the mutations. Then, two PCR fragments were assembled using used the Gibson Assembly kit.

The primers used for the cloning:

5’-TAGGCACCCCACCACCTCAATTGCCAATTAAAAAGGGTCCAGGTGCTCCGCCGGGCAGTA-3’ (F)

5’-GTCGTGGTGGGGCGCCACGACCCCG-3’ (R)

5’-CGGGGTCGTGGCGCCCCACCACGAC-3’ (F)

5’-ATTGAGGTGGTGGGGTGCCTAATGCATTCGGTGGAGCGCCGCGCTTGACT-3’ (R)

Transection was performed in a 10-cm dish at 70%-80% confluence with 30 ul of TransIT 2020 transfection reagent (Mirus Bio, MIR 5400A) and 10μg of total DNA (1µg of pMT-ROWx2FLAG and 9 µg of bluescript plasmids). 12 hours after transection copper (Cu) induction was performed using 500μM of copper. The cells were collected 24h after the induction.

### Co-immunoprecipitation (Co-IP) assay

For each sample, 100µl of fly heads/∼10^7^ S2 cells were homogenized in 500µl/1ml lysis buffer (50mM TRIS HCL 7.4, 150mM NaCl, 1mM EDTA, 1% TRITON X100 and protease inhibitor tablets [Roche]). Homogenate was kept on ice for 30 min and then sonicated using a Bioruptor sonication device (Diagenode) for 2 min (10 sec on 10 sec off). Sonicated lysate was centrifuged at 21,000g for 10 min, and the supernatant was then collected; 35µl were removed for input. Pre-clean: protein A/G PLUS- Agarose beads (Santa Cruz) were washed 3 times with 150µl of lysis buffer, resuspended with 100µl lysis buffer, and then added to the lysate. The samples were rotated for 30 min at 4°C and centrifuged at 3000 rpm for 1 min. The lysate was collected to a new tube and the beads were discarded. IP: 25µl of Red ANTI-FLAG M2 Affinity Gel from SIGMA ALDRICH (F2426) were washed 3 times with TBS (50mM TRIS pH 7.5,150mM NaCl and protease inhibitor tablets [Roche]) and then added to samples. The samples were rotated for 3 hours at 4°C. Next, the samples were centrifuged at 3000 rpm for 1 min and the unbound lysate was collected to a new tube. Beads were washed 3 times using 400µl washing buffer (300mM NaCl, 0.1% NP 40,50mM TRIS HC1 7.5 and protease inhibitor tablets [Roche]) and twice using 400µl of TBS. For western blot, the beads were eluted with SDS-PAGE X1 sample buffer and boiled at 95°C for 5 min.

### Mass spectrometry (MS)

Sample preparation: The packed beads were resuspended in 100μl 8M urea, 10mM DTT, 25 mM Tris-HCl pH 8.0 and incubated for 30 min at 22°C. Next, Iodoacetamide (55mM) was added, and beads were incubated for 30 min (22°C, in the dark), followed by the addition of DTT (20mM). The Urea was diluted by the addition of 6 volumes of 25mM Tris-HCl pH 8.0. Trypsin was added (0.3μg/ sample), and the beads were incubated overnight at 37°C with gentle agitation. The beads were spun down, and the peptides were desalted on C18 Stage tips. Two-thirds of the eluted peptides were used for MS analysis. NanoLC-MS/MS analysis: MS analysis was performed using a Q Exactive Plus mass spectrometer (Thermo Fisher Scientific, Waltham, MA USA) coupled online to a nanoflow HPLC instrument, Ultimate 3000 Dionex (Thermo Fisher Scientific, Waltham, MA USA). Peptides were separated over an acetonitrile gradient (3% to 32% for 45 min.; 32 to 50% for 15 min., 50 to 80% for 10 min.) run at a flow rate of 0.3μl/min on a reverse-phase 25-cm-long C18 column (75μm ID, 2μm, 100Å, Thermo PepMapRSLC). The survey scans (380–2,000 m/z, target value 3E6 charges, maximum ion injection times 50ms) were acquired and followed by higher-energy collisional dissociation (HCD) based fragmentation (normalized collision energy 25). A resolution of 70,000 was used for survey scans. The MS/MS scans were acquired at a resolution of 17,500 (target value 1E5 charges, maximum ion injection times 120ms). Dynamic exclusion was 60 sec. Data were obtained using Xcalibur software (Thermo Scientific). To avoid carryover, the column was washed with 80% acetonitrile, 0.1% formic acid for 25 min between samples. MS data analysis: Mass spectra data were processed using the MaxQuant computational platform (48) (version 1.5.3.12). Peak lists were searched against the UniProt Fasta database of *Drosophila melanogaster*, using both annotated and predicted sequences. The search included cysteine carbamidomethylation as a fixed modification as well as oxidation of methionine as variable modifications and allowed up to two miscleavages. The “match-between-runs” option was used. Peptides with a length of at least seven amino acids were considered, and the required FDR was set to 1% at the peptide and protein level. Protein identification required at least two unique or razor peptides per protein. Relative protein quantification in MaxQuant was performed using the label-free quantification (LFQ) algorithm and with intensity-based absolute quantification (IBAQ). Protein contaminants and proteins identified by less than two peptides were excluded. Statistical analysis between control (n=3) and rowTAG samples (n=3) was performed using the Perseus statistical package(49) (computational platform for comprehensive analysis of (prote) omics data). LFQ values were used as the input for Perseus analysis. Ribosomal proteins were excluded from the results.

### Tagging *row* endogenously by CRISPR/Cas9

Tagging *row* endogenously in *Drosophila melanogaster* using the CRISPR/Cas9 method was performed based on the approach previously described (50) with some modifications. Two gRNAs (one targeting *row* and one targeting the *white* gene) was cloned into pCFD4d plasmid (Addgene plasmid #83954) (as described in the protocol “cloning two gRNAs into plasmid pCFD4”; http://www.crisprflydesign.org/) - (pCFD4d*w/row*)

Primers used for the cloning are: 5’-TATATAGGAAAGATATCCGGGTGAACTTCC CGTGGGGCTTGTATCATTGGTTTTAGAGCTAGAAATAGCAAG-3’ (forward)

5’-CCAAAGAGCAGGAATGGTATATTTTAACTTGCTATTTCTAGCTCTAAAACCCAAAG AGCAGGAATGGTATCGACGTTAAATTGAAAATAGGTC-3’ (reverse).

To generate the donor plasmid for homologous recombination (HR) of the gene *row*, pUC57-white [coffee] plasmid (Addgene 84006) was digested with SacI and HindIII to exclude the donor template for HR of the white gene. We used w^1118^ flies genomic DNA to amplify by PCR both ∼1kb upstream and ∼1kb downstream from the stop codon of the gene *row*. To add the tag, the primers included the FLAG-TAG sequence next to the stop codon. As we used the Gibson Assembly kit, the primers included part of the plasmid backbone as well. The vector backbone and the two parts of the amplified homologous arms were assembled using the Gibson Assembly kit (pUC57- *row*tag).

Primers used for the cloning:

Upstream part: 5’-ACGGCCAGTGAATTCGAGCTCGCGGGTTGAGGTTTATAAGTC-3’ (F)

5’-TCACTTGT*CATCGTCATCCTTGTAAT*CTTGCGGATGGTGATGGTGCT-3’ (R)

Downstream part: 5’-GATTACAAGGATGACGATGACAAGTGATACAAGCCCCAC GGAAA-3’ (F)

5’-CTATGACCATGATTACGCCAGGAGCTATGCCTACCCCTTC -3’ (R)

pCFD4d*w/row*, pUC57-white[coffee], and pUC57-*row*tag plasmids were injected into vas-Cas9 (y1, M{vas-Cas9ZH-2A) flies (50) by Rainbow Transgenic Flies, Inc. Individual injected G_0_ flies were mated with second chromosome balancers flies, and non-red-eyed G_1_-*Cy*O individual flies were crossed to 2nd chromosome balancers flies again. Screening for the desired *row*-FLAG tagged line and elimination of random integration in the G_1_-CyO individuals performed by PCR was as previously described (50). For final validation, we sequenced the entire *row* locus and performed WB as shown in the results section.

### Knockdown experiments in S2 cells

RNAi experiments in S2 cells were performed as previously described(22) with modifications. dsRNA to *row* and *BEAF-32* were generated using the MEGAscript kit (Ambion). Cells were diluted to 10^6^ ml^−1^, and 4 μg of dsRNA per 1ml of cells were added. After two days, the cells were diluted again to 10^6^ ml^−1^, and 8 μg of dsRNA per 1ml of cells were added. After two days, the cells were washed twice with PBS, collected, and used for downstream experiments. The primers used for producing dsRNA to *row* were previously described (22):

Forward *row* T7: TAATACGACTCACTATAGGGTGATACAGACGCTGAGTGATTG

Reverse *row* T7: TAATACGACTCACTATAGGGAGGAACCACATCCCAAGATG

Primers used for producing dsRNA to *BEAF-32*:

Forward BEAF-32 T7: TAATACGACTCACTATAGGGGCGAGGATCCACTGTGCTAT

Reverse BEAF-32 T7: TAATACGACTCACTATAGGGACGCTGATTTGCCCATTTAC

### ChIP-seq using fly heads

The ChIP was performed based on a previous protocol(51) with modifications. 100µl of fly heads were collected using dry ice and homogenized in 1 mL of NEB buffer (10mM HEPES-Na at pH 8.0, 10mM NaCl, 0.1mM EGTA-Na at pH 8.0, 0.5mM EDTA-Na at pH 8.0, 1mM DTT, 0.5% Tergitol NP-10, 0.5mM Spermidine, 0.15mM Spermine and protease inhibitor tablets [Roche]) for a total of 2 min. Homogenate was poured into Bio-Spin chromatography columns (BIO-RAD) and centrifuged at 1000g for 4 min. The filtered homogenate was centrifuged at 6000g for 10 min, and the supernatant was discarded. The pellet was resuspended in 1 mL of NEB and centrifuged at 11,000rpm for 20 min on a sucrose gradient (0.65 mL of 1.6 M sucrose in NEB, 0.35 mL of 0.8 M sucrose in NEB). The Nuclei pellet was then resuspended in 1 mL of NEB, and 11% formaldehyde was added for a final concentration of 1%. Nuclei were crosslinked for 10 min at room temperature and quenched by adding 1/10 vol of 1.375 M glycine. The nuclei were collected by centrifugation at 6000g for 5 min. Nuclei were washed twice in 1 mL of NEB and resuspended in 350mL of Sonication buffer (10mM HEPES-Na at pH 7.5, 2mM EDTA at pH 8.0, 1% SDS, 0.2% Triton X-100, 0.5mM Spermidine, 0.15mM Spermine). Nuclei were sonicated using a Bioruptor sonication device (Diagenode) for 3 cycles of 7 min (30 seconds on 30 seconds off). Sonicated nuclei were centrifuged at 15,000g for 10 min, and the supernatant was frozen at 80°C in 150µl aliquots. 50µl were taken for checking the chromatin quality and the DNA fragment size, which was between 100 to 1000 bp. For each sample, we used 50µl from the aliquot diluted with 500µl of IP buffer (50 mM HEPES/KOH at pH 7.6,2 mM EDTA, 1% Triton, 0.1% Na Deoxycholate in PBS and protease inhibitor tablets [Roche]). Samples were rotated overnight at 4°C after adding antibodies. ChIP-seq was performed with rabbit αHP1c (n=3), αHP1b (n=2), αWOC (n=3), αROW (n=3), αH3K4me3 (n=3) and IgG antibodies (n=2). 20µl per sample of Dynabeads Protein G (Invitrogen) was washed twice with 1ml of IP buffer. Beads were resuspended in 25µl IP buffer per sample. The beads were added to the samples and rotated for 1 hour at 4°C. Chromatin immobilization and barcoding were performed as previously described (52)

### ChIP-seq and ChIP-qPCR using S2 cells

The ChIP was performed based on a previous protocol (22) with modifications. For each sample, a 25 cm flask with 8-10×10^6^ S2 cells was collected into a 15 ml tube and formaldehyde was added to a final concentration of 1.8%. The cells were crosslinked for 10 min at RT on a shaker. To stop the reaction, glycine was added to a final concentration of 125mM (from a stock solution of 2.5M in PBS). The cells were centrifuged for 3 min at 1500g at 4°C, the supernatant was discarded, and the cells were resuspended in 2ml PBS. Then, the cells were centrifuged again (3 min at 1500g at 4°C), the supernatant was discarded, and the cells were resuspended in 1ml washing buffer A (10mM Hepes pH7.9, 10mM EDTA 0.5mM EGTA 0.25% Triton X100m, protease inhibitor). The 1ml suspension was transferred to an Eppendorf, incubated for 10 min at 4°C on a wheel, and then centrifuged (3 min at 1500g at 4°C). The supernatant was discarded, the pellet resuspended in 1ml washing buffer B (10mM Hepes pH7.9, 100mM NaCl, 1mM EDTA, 0.5mM EGTA, 0.01 % Triton X100) and incubated for 10 min at 4°C on a wheel. The suspension was centrifuged again (3 min at 1500g at 4°C), the supernatant was discarded, and the pellet was resuspended in 450µl TE buffer. 50µl 10% SDS was added, and the Eppendorf was inverted 5 times and centrifuged at 1500g at 4°C for 3min. The upper phase was carefully removed with a pipette, 500µl TE was added, and the Eppendorf was inverted 5 times, after which it was centrifuged. The last procedure was repeated once again. After the second centrifuge, the upper phase was removed, TE with 1mM PMSF was added to a final volume of 400µl, and 4 µl of 10% SDS was added. The suspension was sonicated using Bioruptor plus sonication device (Diagenode) for 20 cycles (30 sec on 30 sec off).

The following solutions were added in this order to the lysate, and between each addition, the lysate was incubated on the wheel at 4°C for 2min: 42µl 10% TRITON (final 1%), 4.2µl 10% Deoxycholate (final 0.1%), and 15µl 4M NaCl (final 140 mM). The lysate was Incubated on the wheel at 4°C for 10min and centrifuged at full speed at 4°C for 5min. The supernatant was divided into aliquots and was frozen at -80°C; 45µl were kept for input and for checking DNA fragment size. For the ChIP-seq experiment, 50 µl of chromatin aliquot was used and diluted with 450 RIPA buffer (140 mM NaCl, 10 mM Tris-HCl pH, 1 mM EDTA, 1 % TritonX100, 0.1 % SDS, 0.1 % Deoxycholate and protease inhibitor). Samples were rotated overnight at 4°C after adding antibodies. Washes performed as described in the “ChIP-seq using fly heads” paragraph. Chromatin immobilization and barcoding were performed as previously described (52)

For the ChIP-qPCR 100 µl of chromatin aliquot was used and diluted with 1m of ChIP buffer (10 mM Tris-HCl [pH 8.0], 5 mM EDTA [pH 8.0], 150 mM NaCl, 1% Triton X-100 and protease inhibitor). Anti-FLAG M2 magnetic beads (Sigma) were added and samples were rotated overnight at 4°C. Washes were performed with the ChIP buffer (x4) and with TE (x4).

qPCR was performed with primers to three promoters containing ROW AT-motif and one exon not bound by ROW as a control.

Primers are shown below:

5’-TTTGTGTGTTCCATTGCGTA-3’ (Hcf promoter F)

5’-TGCCCACTTATTTGACCGCA-3 (Hcf promoter R)

5’-CAGTCACGTCACGAGAGCAT-3’ (Hcs promoter F)

5’-GGTCATATCTGGCGGGTTTT-3’ (Hcs promoter R)

5’-TATGTTGTTCACCCCCACCT-3’ (Phax promoter F)

5’-AGCGGCAGAGTGACCATACT-3’ (Phax promoter R)

5’-AATACCGTCAACGGATCAGC-3 (Unbound prl-1 exon F)

5’-CAGTTCCCGTTTTGTTTTCG-3’ (Unbound prl-1 exon R)

### ChIP-seq analysis

ChIP-seq reads were aligned by bowtie2 software to the full genome sequence of *Drosophila melanogaster* (UCSC version dm3). Duplicate reads were removed using the samtools rmdup function, and uniquely mapped reads were kept (samtools view -q 10). Peak calling was performed with the callpeak function of the MACS2 tool (53) with a q-value<0.05. An IgG sample was used as a control. Peaks in ChrU and ChrUextra were excluded for subsequent analysis. For visualization, bedGraph files were created by deepTools (a python tool) with bamCoverage function and normalization parameter set to RPKM. The bedGraph file with the average between replicas was generated, and the corresponding bigWig file was created. The bigWig files were uploaded to the integrative genomic viewer (IGV). The BigWig file of BEAF-32 in Kc167 cells was downloaded from NCBI (GSM762845) (35). The ComputeMatrix and plotProfile functions of deepTools were used for plotting the distribution of the ChIP-seq targets relative to the positions of TSSs or TAD boundaries. For each target, the average across replicates was used. UCSC refGene annotation file was used to determine the TSS locations. Classified TADs locations were obtained from the Chorogenome Navigator (7). The correlation of ChIP-seq signal across the genome was calculated based on non-overlapping bins of 2000bp. The deeptools pyBigWig python extension was used for calculating the average signal in each bin. A pairwise correlation was calculated with the psych R package. The locations of peaks within different genes annotation were determined using the annotatePeak function from the ChIPseeker R package (54). The function enrichPeakOverlap from ChIPseeker was used to test the overlap of ChIP-seq peaks with different histone modifications. Histone modification data were from modENCODE ChIP-chip data (30). Multiple logistic regression implemented in the HiCfeat R package (38) was used to calculate the enrichment and influence of proteins on TAD boundaries. UCSC refGene annotation was used to define the genes bound by ROW or BEAF-32. Promoter regions (between - 250bp and +50 of TSS) of genes that overlapped with ROW/BEAF-32 peaks were considered to be bound by the proteins (Table S2 and Table S4). A BED file of BEAF-32 called peaks in Kc167 cells was downloaded from NCBI (GSM762845) (35).

Motif enrichment analysis was performed using the MEME-ChIP motif analysis tool (55) with the default parameters. Input sequences were taken from an interval that included 250bp downstream and 250bp upstream of the peak’s summit. CentriMo option (56) with default parameters was used to calculate the probability to find the motif relative to the peak summit. Genes bound by ROW were defined as direct targets if at least 15 A or T repeats were observed at their promoter and as indirect targets if the motif was not found. Enrichment of GAF, NELF-E, and CP190 with genes unbound, bound directly and indirectly by ROW was calculated using Fisher’s exact test. Data was downloaded from GSE40646 (40) (GAF), GSE116883 (57) (NELF-E), and GSE41354 (58) (CP190). Gene ontology (GO) enrichment analysis for genes bound by ROW was performed using the FlyEnrichr tool (36, 37). Redundant GO terms were removed using the REVIGO tool (59).

### Predicting DNA-binding specificities for Cys2His2 zinc fingers

ROW protein sequence (download from UCSC) was used as input in the online tool for predicting zinc finger DNA binding (http://zf.princeton.edu/logoMain.php) (34).

### MNase-seq

100µl of fly heads per sample was homogenized for 5 min in a homogenization chamber with 300µl of ice-cold Nuc. Buffer-I (60mM KCl, 15mM NaCl, 5mM MgCl_2_, 0.1mM EGTA, 15mM Tris-HCl pH=7.5, 0.5mM DTT, 0.1mM PMSF and 1x complete protease inhibitor (Roche)). Next, 200µl more of Nuc. Buffer-I was added. The homogenate liquid was poured into Bio-Spin chromatography columns (BIO-RAD) and centrifuged at 1000g for 1 min at 4°C. 500µl of ice-cold Nuc. Buffer-II (0.04% NP-40 and Nuc. buffer I) was added to the filtered homogenate, mixed by inverting the tube, and incubated for exactly 7 min on ice. During the 7 min incubation, 13ml tubes with 8ml ice-cold Nuc. Buffer-III (1.2M sucrose + Nuc. buffer I) was prepared. The 1ml filtered homogenate was poured into the 13 ml tube of Nuc. Buffer- III. The tube was centrifuged for 10 min at 11,000rpm. The supernatant was carefully removed so that only the pellet remained (containing the intact purified nuclei). The nuclei pellet was re-suspend in 500µl ice-cold MNase digestion buffer (0.32M sucrose, 50mM Tris-HCl pH=7.5, 4mM MgCl_2_, 1mM CaCl_2_ and 0.1mM PMSF). The 500µl nuclei were split into 2 Eppendorf with 250μl in each. 5 units (50 Gel Units) of Micrococcal Nuclease (NEB) were added to one tube and 20 units (200 Gel Units) to the other tube (for over digesting). The tubes were inverted for mixing and incubated at 37°C for 10 min. The reaction was stopped by the addition of 13μl of 20mM EDTA and 13μl of 10% SDS in each tube, followed by inverting and cooling on ice. 2.8μl of 10mg/ml Proteinase K was added and mixed by inverting. The nuclei were incubated at 65°C overnight and then centrifuged for 1 min at 15,000g. The supernatant from each tube was taken and cleaned separately by PCR clean-up kit (Invitrogen) with 26µl of elution buffer at the final step. Next, 1µl of 0.3ug/µl RNase free DNase (Sigma), 0.75μl of 0.5M Tris-HCl pH=7.4, and 3.5μl 120mM NaCl were added to each tube and mixed by pipetting and spin-down. The tubes were then incubated for 1h at 37°C, and the mix was cleaned again using a PCR clean-up kit (Invitrogen) with 12µl of elution buffer at the final step. Each tube volume was put to run on a 1.5% agarose gel alongside a 100bp DNA ladder. Mononucleosome bands (150bp-200bp) from the gel were excised and cleaned with a gel clean-up kit (Invitrogen) with 50µl of DDW as a final step. The purified DNA was quantified by nanodrop, and an equal amount from the two digestion levels was taken for DNA library preparations. DNA libraries were made as previously described (60).

### RNA purification and libraries preparation for RNA-seq using fly heads

We crossed *Actin GAL4*/CyO flies with *row*^RNAi-1/2^ flies or with w^1118^ control flies and selected 0-3 days old *row*^RNAi-1^ (*Act-GAL4*/+; UAS-*row*^RNAi-1^/+), *row*^RNAi-2^ (*Act-GAL4*/+; UAS- *row*^RNAi-2^), and *Act-GAL4* Control (*Act-GAL4* /+) flies. The flies were raised for three more days to recover from CO_2_ exposure and collected using dry ice. RNA was prepared using TRI Reagent (SIGMA) according to the manufacturer’s protocol. 3’ RNA-seq library was performed based on Guttman’s lab “RNAtag Seq protocol” (61) with modifications. Fragmentation of RNA was performed using FastAP Thermosensitive Alkaline Phosphatase buffer (Thermo Scientific) for 3 min at 94°C. Samples were placed on ice, and then the FastAP enzyme was added for 30 min at 37°C. RNA was purified using 2.5× volume of SPRI beads (Agencourt), and then linker ligation was performed with an internal sample-specific barcode using T4 RNA ligase I (NEB). All RNA samples were pooled into a single Eppendorf and purified using RNA Clean & Concentrator columns (Zymo Research). Poly A selection was performed with Dynabeads Oligo(dt) beads according to the manufacture’s protocol. RT was performed with a specific primer (5′-CCTACACGACGCTCTTCC-3′) using AffinityScript Multiple Temperature cDNA Synthesis Kit (Agilent Technologies). RNA degradation was performed by incubating the RT mixture with 10% 1 M NaOH (2μl of RT mixture, 70°C, 12 min). The pH was then neutralized by AcOH (4 μl for 22 μl mixture). Next, the cDNA was cleaned using Silane beads (Life Technologies). The 3′-end of the cDNA was ligated to linker2 using T4 RNA ligase I. The sequences of linkers are partially complementary to the standard Illumina read1 and read2/barcode adapters, respectively. The cDNA was cleaned using Silane beads, and PCR was performed using enrichment primers and Phusion HF Master Mix (NEB) for 12 cycles. The library was cleaned with 0.8× volume of SPRI beads.

### RNA purification and libraries preparation for RNA-seq using S2 cells

RNA purification was performed with RNeasy Mini Kit (Qiagen) according to the manufacturer’s protocol. 3’ RNA-seq library generation was performed based on a previous protocol(62). In brief, 100ng from each sample was incubated with oligo-dT reverse transcription (RT) primers containing 7bp barcode and 8bp UMI (Unique Molecular Identifier) at 72°C for 3 minutes and moved directly to ice. This was followed by a Reverse transcription reaction (SmartScribe kit Clontech) for 1 hour at 42°C and 70°C for 15 minutes. Next, samples were pooled and purified using SPRI beads X1.2 (Agencourt AMPure XP, Beckman Coulter). The pooled barcoded samples were then tagmented using Tn5 transposase (loaded with oligos Tn5MEDS-A) for 8 min at 55°C Followed by the addition of 0.2% SDS to strip off the Tn5, and SPRI X2 purification was performed. Finally, a PCR was performed with primers containing NGS sequences (KAPA HiFi HotStart ReadyMix, Kapa Biosystems, 12 cycles), and the library was cleaned using 0.8x SPRI beads.

### Analysis of RNA-seq data

RNA-seq reads were aligned by STAR (63) to the full genome sequences of *Drosophila melanogaster* (UCSC version dm3) and counted per annotated gene using ESET (End Sequencing analysis Toolkit) (64). Differential expression analysis was performed using the edgeR R package. In the data from fly heads, we compared the two *row*^RNAi^ lines to the control (65). Only genes with a count per million (CPM) > 1 in at least 3 different samples were included in the analysis. Differentially expressed genes were selected by FDR<0.05, with no fold-change cutoff. The ranked list of differentially expressed genes based on FDR (Table S1) was used as an input for the enriched GO terms tool – Gorilla (66). The REVIGO tool (59) was used to remove redundant GO terms. In the data from S2 cells, we compared all treated samples to untreated samples. The FilterByExpr function in the edgeR R package was used to filter out non-expressed genes. Differentially expressed genes were selected by FDR<0.1, with no fold-change cutoff (Table S5). The associations between genes that were significantly differentially expressed with other lists of genes (For example genes bound by ROW or BEAF-32 and BEAF-32 long-range targets) were tested using Fisher’s exact test.

The gene expression variation for genes bound or unbound by ROW was calculated using published *Drosophila* gene expression data from 30 developmental stages (67). Correction of the variation to expression levels was performed by calculating the residuals from a loess curve (locally estimated scatterplot smoothing) fitted to scatterplot between the average genes expression (log2) and the coefficient of variation (log2) in 30 developmental stages. The residuals, which are not correlated with the expression levels, were presented as a corrected variation for genes bound and unbound by ROW.

### Analysis of Hi-C data

Hi-C data was downloaded from NCBI: GSE97965 (7) and processed using HiCExplorer (7, 68). Each mate of paired-end was aligned separately with bowtie2 to the full genome sequence of *Drosophila melanogaster* (UCSC version dm3) with local and reorder parameters. The contact matrix was created using the hicBuildMatrix function of the HiCExplorer tool with a bin size of 1kb and corrected using the hicCorrectMatrix function with Knight-Ruiz (KR) method parameter.

To create the aggregation plots hicAggregateContacts function of the HiCExplorer tool was used. In the bed parameter, a bed file with the locations of the direct peaks of ROW was used, and in the BED2 parameter, a bed file with the locations of the indirect peaks of ROW was used. In the OperationType parameter, we chose the mean, and in the transform parameter, we chose obs/exp. The range was set to 10kb-500kb. As a control, the same number of promotors not bound by ROW were selected randomly and used as input in the hicAggregateContacts function with the same parameters.

To identify over-represented promoter interactions, we used the Hi-C analysis tool PSYCHIC (41). A symmetric matrix for each chromosome in 5kb resolution was created and used as input. UCSC refGene annotation file was used as the genes file input. We used the output bed file of over-represented pairs with FDR value < 0.05. The file contains genes and putative long-range interactions location. Using Fisher’s exact test, we calculated the association between genes with a direct ROW peak (with AT motif) at the promoter region and putative long-range interactions locations with an indirect ROW peak (without AT motif). Output file of random interactions with genes was used as control.

## Acknowledgments

This work was supported by The Legacy Heritage Bio-Medical Program of the Israel Science Foundation [Grant No. 839/10 to SS and SK] and by the Israel Science Foundation (grant no. 575/17 to SS).

## Author contributions

NH performed the experiments, analyzed the data, and wrote the manuscript. SS and KS conceived the study, supervised the project and wrote the manuscript.

## Availability of data and materials

The datasets supporting the conclusions of this article are included within the article and its additional files or are available in the Gene Expression Omnibus database under accession number GSE169415 https://www.ncbi.nlm.nih.gov/geo/query/acc.cgi?acc=GSE169415.

