## Supplementary Figures for "The chromatin factor ROW cooperates with BEAF-32 in regulating long-range inducible genes"

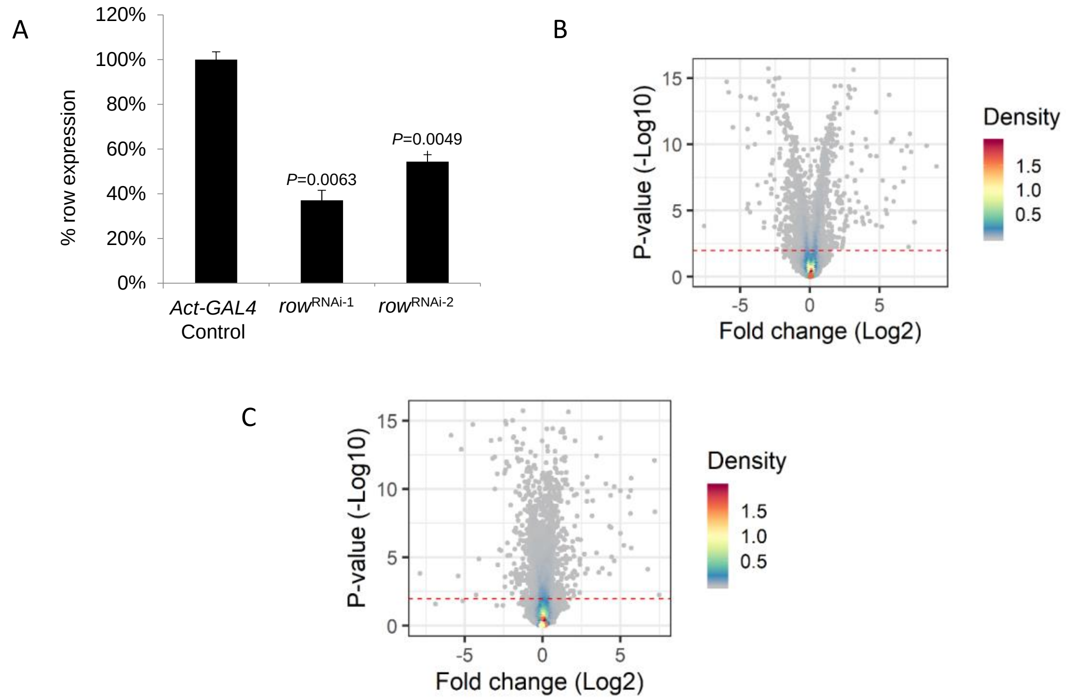


**Figure S1**

(A) Expression of *row* is significantly decreased in *row*$^{RNAi\text{-}1}$ and *row*$^{RNAi\text{-}2}$ flies based on RNA-Seq results.

(B-C) Volcano plot of the *P*-values (-log10) as a function of the gene expression fold change (log2) in (B) *row*$^{RNAi\text{-}1}$ and (C) *row*$^{RNAi\text{-}2}$ relative to *Act-GAL4* control flies.

A

B

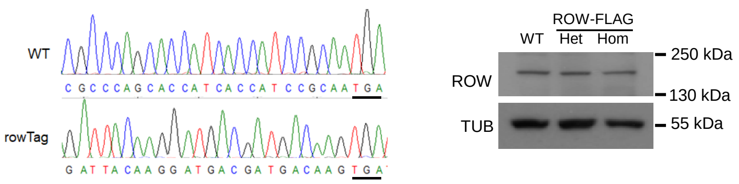


C

| Protein names | Control IBAQ | ROW-TAG IBAQ | Peptides number |
|---|---|---|---|
| HP1c | 0 | 2.44E+08 | 9.3 |
| HP1b | 0 | 1.69E+08 | 10.7 |
| ROW | 0 | 7.93E+07 | 47.7 |
| WOC | 0 | 7.73E+07 | 62.3 |
| Ctp | 0 | 3.06E+07 | 2.0 |
| Ubqn | 0 | 2.54E+07 | 10.7 |
| SsRbeta | 0 | 3.90E+06 | 0.7 |
| TM4SF | 0 | 3.13E+06 | 1.0 |
| CG16985 | 0 | 3.04E+06 | 1.0 |
| BEAF-32 | 0 | 2.73E+06 | 2.7 |
| Sgs3 | 0 | 2.62E+06 | 1.0 |
| Rab1 | 0 | 2.05E+06 | 1.7 |
| CathD | 0 | 1.82E+06 | 0.7 |
| CG9281 | 0 | 1.69E+06 | 1.0 |
| Adh | 0 | 1.26E+06 | 0.7 |
| SdhC | 0 | 1.15E+06 | 0.7 |
| CG3192 | 0 | 1.01E+06 | 0.7 |
| Trip1 | 0 | 8.78E+05 | 1.3 |
| CG33303 | 0 | 7.57E+05 | 1.0 |
| CG6126 | 0 | 6.71E+05 | 0.7 |
| Lsp1gamma | 0 | 6.42E+05 | 1.7 |
| CG4045 | 0 | 6.20E+05 | 1.0 |
| CG5973 | 0 | 5.00E+05 | 0.7 |
| Rack1 | 0 | 4.79E+05 | 0.7 |
| CG1737 | 0 | 4.50E+05 | 2.7 |
| Whd | 0 | 3.27E+05 | 1.3 |
| gammaCop | 0 | 2.68E+05 | 0.7 |
| CG32638 | 0 | 2.40E+05 | 0.7 |
| Chro | 0 | 2.21E+05 | 1.7 |
| Pst | 0 | 2.15E+05 | 1.0 |
| Tspo | 0 | 2.14E+05 | 0.7 |
| CASK | 0 | 1.96E+05 | 0.7 |
| T-cp1 | 0 | 1.90E+05 | 0.7 |
| Foxo | 0 | 1.48E+05 | 1.0 |
| Flacc | 0 | 1.33E+05 | 0.7 |
| Hfp | 0 | 1.30E+05 | 0.7 |
| E(bx) | 0 | 8.37E+04 | 0.7 |
| Crol | 0 | 4.16E+04 | 0.7 |

D

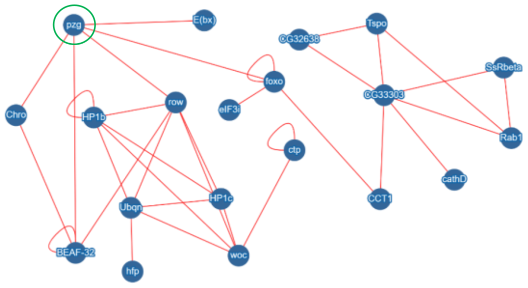


**Figure S2**

(A) Sanger sequencing results upstream to the stop codon (underlined) of the gene *row* from WT and endogenously ROW-FLAG tagged flies.

(B) Western blot for flies expressing endogenous FLAG-tagged ROW (ROW-TAG) with αROW antibody used to verify that there is no effect of the tagging on ROW protein levels. αTubulin (TUB) antibody was used as a reference. Het, Heterozygotes; Hom, Homozygotes for tagged ROW.

(C) List of proteins co-purified with ROW in at least two out of the three experiments and were not detected in the control experiments. IBAQ (Accurate Label-Free Protein Quantitation) reflects the protein abundance in the sample. Peptides number is the number of Razor and unique peptides. The data are the mean of control (W1118, n=3) and ROW-FLAG flies (n=3) samples.

(D) Proteins-protein interaction supported by previous studies based on the molecular interaction search tool (MIST) [1]. The PZG (Z4) protein (circled in green) was not co-purified with ROW in our study but was found previously to interact with ROW [2,3] and may connect the two complexes co-purified with ROW in our experiment.

A

| ROW Sequence | E-value |
|---|---|
| TWWWTATTTWTWWTWTWWWTW | 1.50E-254 |
| GAATTAAATAT | 3.90E-115 |
| ATATWTATWATWRT | 1.80E-114 |
| AAATRTW | 8.40E-82 |
| TTAATTA | 6.50E-81 |

| WOC Sequence | E-value |
|---|---|
| WTWTATTTATWWWWTTWTTTW | 1.30E-172 |
| GAATTAAATAT | 9.90E-69 |
| ATATWTATWATWRT | 1.10E-65 |
| AATAAW | 6.20E-44 |
| AAATAT | 1.10E-43 |

| HP1c Sequence | E-value |
|---|---|
| TWWWTWTWTTTWTTWWTTTWTTTTWTTTTT | 2.20E-175 |
| TAAWAATA | 5.00E-21 |
| GAATTAAATAT | 5.20E-19 |
| ATATWTATWATWRT | 4.40E-17 |
| MATAAAAAAAA | 9.00E-16 |

| HP1b Sequence | E-value |
|---|---|
| AAAAAAWAAAAHAAAAAAAWAAAWAAAA | 1.30E-146 |
| WAAATATW | 1.50E-21 |
| ATATWTATWATWRT | 8.30E-15 |
| GAATTAAATAT | 6.10E-09 |
| TAAWAATA | 1.10E-08 |

B

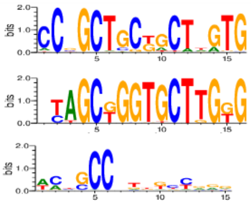

C

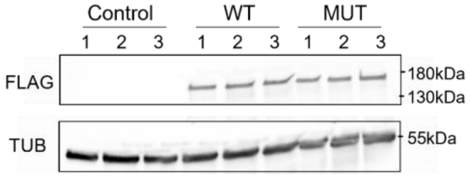


**Figure S3**

(A) Top five centrally enriched sequences in ROW, WOC, HP1b, and HP1c binding sites in fly heads based on MEME-ChIP tool.

(B) Predicted DNA sequences for ROW- Cys2His2 zinc fingers under different models.

(C) Validation of the expression of ROW-FLAG tagged proteins in S2 cell transfected with WT or AT-hook mutant ROW-FLAG tagged plasmids (MUT) by western blot. Control are cells not transfected with ROW-FLAG tagged plasmids. Western blot was performed with αFlag tag and αTubulin (TUB) antibodies.

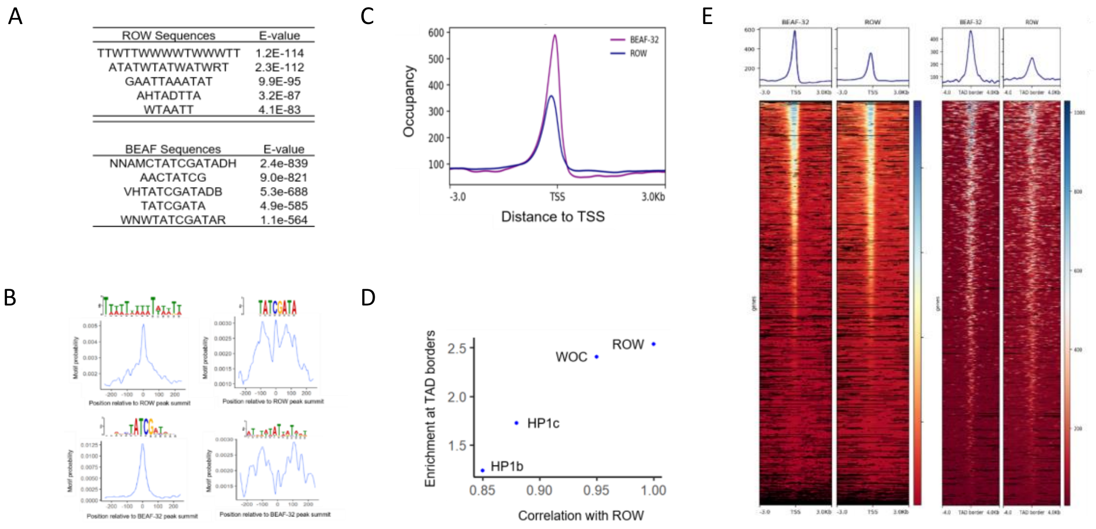


**Figure S4**

(A) Top five centrally enriched sequences in ROW and BEAF-32 binding sites in S2 cells based on MEME-ChIP tool.

(B) Central enrichment of AT-rich sequences and BEAF-32 consensus motifs (logo) in ROW binding regions (upper figures) and BEAF-32 binding regions in S2 cells (lower figures). The plot shows the probability of having the motif relative to ROW or BEAF-32 ChIP-seq peak's summits (calculated with centriMo).

(C) The distribution of ROW and BEAF-32 binding relative to the positions of TSSs in S2 cells.

(D) Binding enrichment at TAD boundaries for ROW, WOC, HP1c, and HP1b ($\hat{\beta}$) as a function of the correlation of ChIP-Seq signals for each protein with ROW in fly heads.

(E) Heatmap of ROW and BEAF-32 ChIP-Seq signals in S2 cells ± 3 kb around TSS (left figure) or ± 4 kb around TAD boundaries (right figure).

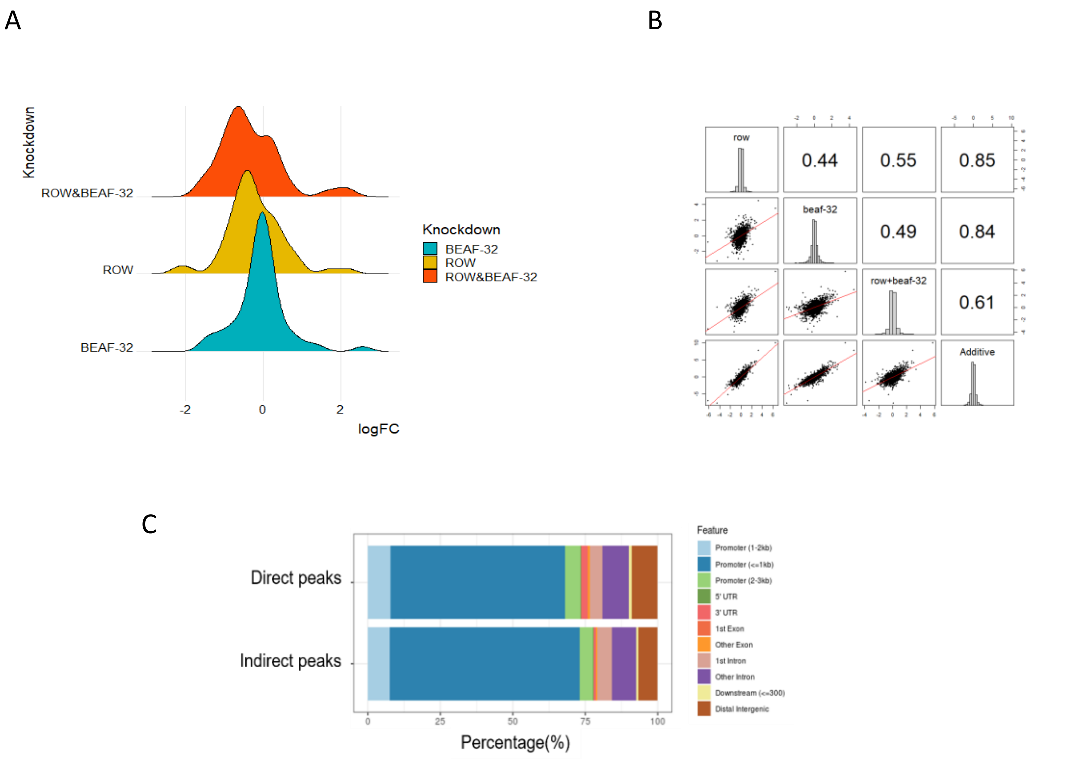


**Figure S5**

(A) Density plot of the gene expression fold changes (log2) in cells treated with rowdsRNA, BEAF-32dsRNA, and rowdsRNA + BEAF-32dsRNA, shown specifically for significantly differentially expressed genes.

(B) Pairwise correlation between the gene expression fold changes (log2) in cells treated with rowdsRNA, BEAF-32dsRNA, rowdsRNA + BEAF-32dsRNA, and additive model (sum of fold change in the separate knockdowns of row and BEAF-32). Fold changes were calculated relative to control cells (using edgeR). In the middle: the distributions of the fold-changes. On the bottom: bivariate scatter plots with linear regression lines are displayed. On the top: the value of the correlation.

(C) Genomic annotation of direct and indirect peaks of ROW show that most are gene promoters.
